# The Phenotypic Landscape of a Circadian Clock

**DOI:** 10.64898/2026.04.23.720472

**Authors:** Soo Ji Kim, Diane Schnitkey, Bryan Andrews, Rama Ranganathan, Michael J. Rust

## Abstract

Circadian clocks produce near-24-hour oscillations through biochemical feedback loops. To study their architecture, we developed a deep sequencing assay that measures the phenotypes of thousands of mutant clocks in parallel. We reveal a landscape where oscillator properties are factorized: mutations change period without decreasing amplitude and while maintaining a balanced waveform. Mutations that either shorten or lengthen period localize to specific protein-protein interaction surfaces, while a particularly sensitive region near the KaiC interdomain linker can cause extreme effects. After entrainment, high amplitude mutant oscillators form a tunable low-dimensional manifold in the period-phase plane, suggesting that most period mutations leave the coupling to the environment unchanged. In contrast, mutations that reduce amplitude are concentrated in a specific long period phenotype. This correlation structure may support the evolvability of this dynamical molecular system and is a powerful constraint on underlying mechanism.

## Main Text

Much has been learned about the architecture of proteins using deep mutational scanning approaches, which enable the analysis of all possible single amino acid substitutions (*1, 2*). However, cellular physiology is not generally reducible to the action of isolated proteins. Rather, networks of interacting components emergently produce complex dynamic phenotypes including environmental sensing, cell cycle control, cell-to-cell signaling, and internal oscillations. Thus, an urgent problem is to understand the phenotypic landscape of such complex traits in macromolecular systems under mutation and the resulting consequences for evolution.

The cyanobacterial circadian clock represents a uniquely powerful window into this problem because it generates complex dynamical behavior from a small module of genes. Cyanobacteria exhibit near-24 hour oscillations in gene expression that have all the hallmarks of the circadian rhythms seen in multicellular organisms: they maintain high amplitude rhythms in a constant environment, the rhythms entrain to light-dark cycles in the environment, and the period of oscillation is temperature compensated (*3*). Remarkably, oscillations can be reconstituted *in vitro* using purified proteins KaiA, KaiB, and KaiC which self-organize to create a coherent 24-hour rhythm in KaiC phosphorylation (*4*). In vivo, the *kaiBC* promoter is a direct target of output signaling so that the abundance of the *kaiBC* mRNA oscillates during the circadian cycle (*5*).

Post-translational oscillations in the KaiABC network are sustained by a negative feedback loop. KaiC is a hexameric double-domain ATPase that is progressively autophosphorylated on its C-terminal (CII) domain throughout the course of the day (*6, 7*). Phosphorylation is stimulated by KaiA (*8*), which binds to the tail of CII (*9*). Near the end of the day, CII phosphorylation triggers allosteric signaling across KaiC to the N-terminal ATPase domain (CI) (*10, 11*). KaiB then binds to CI and forms an inhibitory ternary complex that traps and inactivates KaiA (*6, 12*), closing the negative feedback loop. With KaiA inactive, KaiC dephosphorylates, allowing the cycle to repeat (*13*). Transcriptional output is achieved by rhythmic docking of histidine kinases (SasA and CikA) onto the KaiC complex, which control activation of an output transcription factor RpaA that binds directly to the *kaiBC* promoter and many others (*5, 12, 14*).

Multiple factors make the *kaiABC* gene cluster an appealing target to map out a multidimensional phenotypic landscape. First, the sufficiency of the KaiABC proteins *in vitro* suggest that the system can be treated as a self-contained oscillator module. Second, individual cyanobacterial cells maintain coherent rhythms for many cycles, and there is negligible coupling between cells (*15, 16*). Third, the natural competence of cyanobacteria allows us to stably integrate large barcoded libraries of gene variants into the chromosome—thus we can study the impact of genetic variation on the output of a nonlinear dynamical system in its natural organismal context (*17*).

### Assaying thousands of KaiABC variants in parallel

We envisioned a strategy to study a library of mutant circadian clocks coexisting in a single culture by measuring the dynamics of each mutant in parallel. To read out each mutant phenotype, we exploited the rhythmic expression of the *kaiBC* transcript: each mRNA carries an untranslated barcode which we reverse transcribe and sequence over a multiday time course. Thus, the temporal dynamics of each RNA barcode reports on its own mutant phenotype (Fig. 1A).

**Figure 1.**
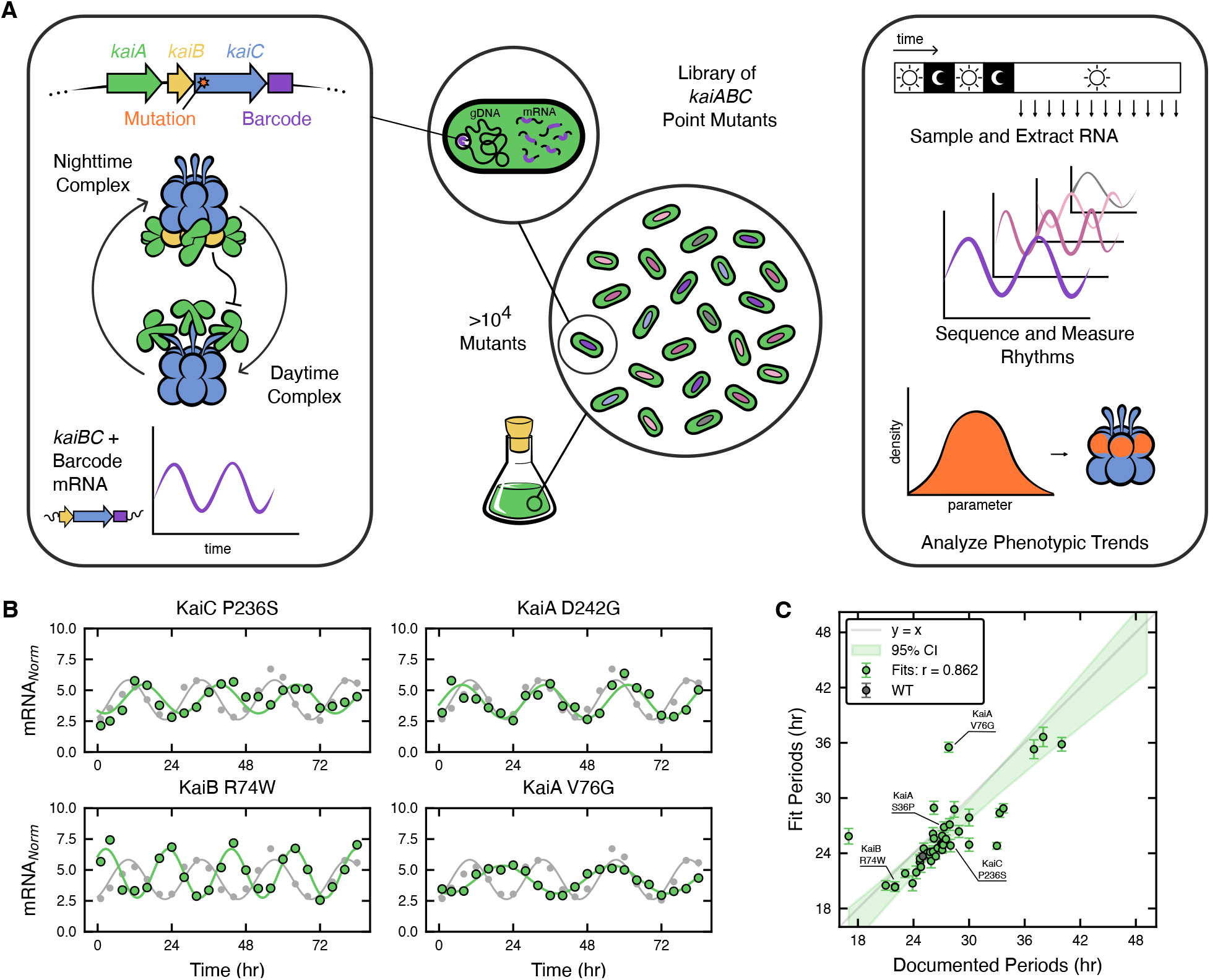
Deep mutational scan of a circadian clock. (**A**) In *Synechococcus elongatus* the KaiABC proteins create a circadian rhythm. The proteins switch between two alternative complexes: in the daytime complex, KaiA binds to the tail of the KaiC C-terminal domain (CII) and promotes KaiC autophosphorylation. In the nighttime complex, KaiC phosphorylation triggers binding of KaiB to the KaiC N-terminal domain (CI) which inhibits KaiA through sequestration, forming a negative feedback loop. The output of this post-translational cycle generates transcriptional rhythms including in the *kaiBC* transcript. We generated a pooled library of mutations in the *kaiABC* genes, barcoded in the 3’-UTR (see Fig. S1), then extracted RNA every 4 hours under constant light. The dynamic phenotype of each mutant can be identified by the time course of each RNA barcode (see Fig. S2) and resulting phenotypic properties localized to the protein structures. (**B**) Rhythms of previously known period mutants (green) compared to the WT rhythm (gray). Barcode read counts are normalized to represent RNA expression per genome relative to expression from arrhythmic nonsense mutants. (**C**) Comparison of oscillator periods from previously reported mutants (*green*) and WT (*black*) with their phenotypes in the deep mutational scan. Error bars represent uncertainty in the least-squares fit for period. The 95% confidence interval is calculated from linear regression with a slope of 0.862. *r* is the Pearson correlation coefficient.

Following this strategy, we created a deep mutational scanning library in the *kaiABC* gene cluster where over 11,000 single amino acid substitutions in the proteins are represented at high read depth (Fig. S1). After an initial synchronization with light-dark cycles, we allowed the mutant oscillators to free run and collected an RNA timecourse over the following 84 hours. Barcode reads mapping to each amino acid substitution were pooled and normalized by DNA abundance and by the expression level of barcodes mapping to premature STOP codons in KaiC. This normalization creates an output signal indicating the expression of *kaiBC* per cell relative to a true null phenotype. We then used a sinusoidal least-squares regression approach to extract estimates of amplitude, period, phase, and baseline for each mutant (Figs. S2, S3, Data S1, S2). We compared the barcode signal of wildtype and known period mutants, observing a clear change in frequency caused by mutation (Fig. 1B). Previously reported period mutants in our library correlate well with their literature values (Fig. 1C) (*3, 18-20*).

### A multimodal distribution of amplitudes

The distribution of mutant phenotypes revealed in this assay shows a multimodal amplitude structure that can be decomposed using a Gaussian mixture model (Fig. 2A). The mean of the high amplitude peak is similar to the WT amplitude, while a small very low amplitude shoulder primarily represents complete loss of function in KaiA and KaiC and overlaps with phenotypes caused by premature stop codons. The intermediate peak retains dynamic transcriptional output, but has either low amplitude or decaying oscillations relative to wildtype (Fig 2B, S4) and has a depressed baseline (Fig. S3).

**Figure 2.**
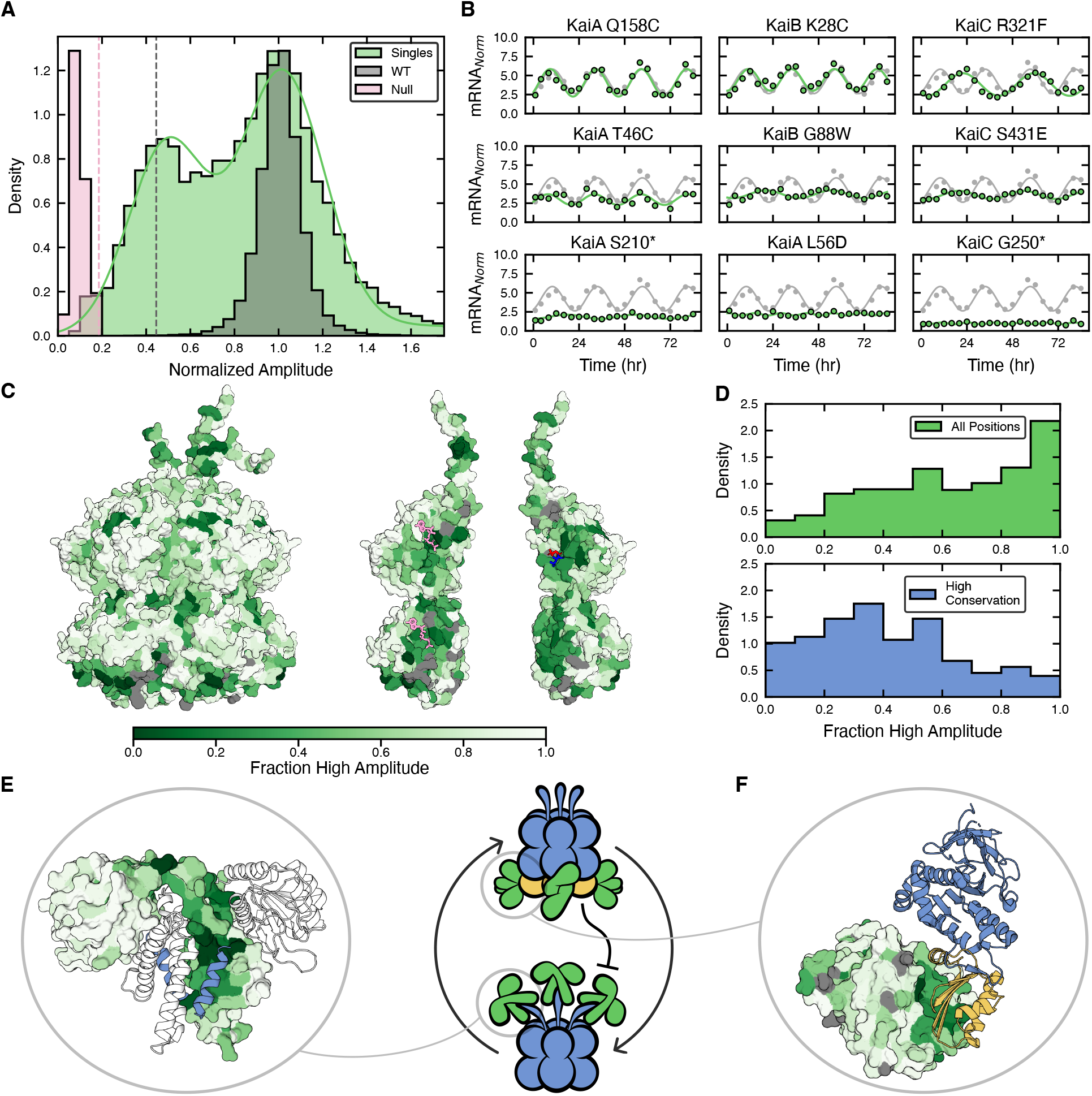
A multimodal amplitude structure linked to conservation, catalysis, and KaiA-dependent feedback. (**A**) Distribution of amplitude normalized to WT. Gray shows amplitude for all sufficiently read single mutants in the library (n = 9,445). This distribution was fit to a gaussian mixture model, defining a high amplitude and low amplitude mode. The null normalization set distribution (*pink*) contains very low amplitude signals coming from missense mutations (n = 71). The WT distribution (*green*) consists of WT barcodes that were subsampled to have similar read depth to typical point mutations. (See Fig. S2). Each distribution is rescaled to have the same peak height. (**B**) Examples of single mutants belonging to different amplitude groups. Top row: WT-like high amplitude. Middle row: low amplitude. Bottom row: null group. (**C**) KaiC structure where each position is colored by amplitude score: the fraction of mutations that cause low amplitude. Left: KaiC hexamer structure (3DVL) (*21*). Right: cut-away views of the KaiC subunit interfaces. ATP molecules located in subunit interface are shown in pink. The phosphorylation sites at 431 and 432 are colored blue and red. Residues with fewer than 5 mutants are colored gray. (**D**) Fraction of mutations causing high amplitude for all single mutants (*gray*) and mutations at highly conserved residues (*green*). High conservation is defined as the positional KL divergence > 2.5. (See SI methods). (**E**) One subunit of the KaiA dimer (5C5E) (*47*) colored by the high amplitude fraction of each residue. The other subunit is shown in ribbon representation. Blue helix is the C-terminal fragment of KaiC bound to KaiA (*48*) (**F**) Structure of the KaiABC nighttime complex (5JWR) (*12*). The surface of the KaiA C-terminal domain is colored by high amplitude fraction. Blue ribbon: KaiC CI domain, yellow ribbon: KaiB.

Mutations at the KaiC subunit interfaces, around the catalytic sites, and within the hydrophobic core of the protein tend to cause loss of amplitude, but not a null phenotype, and are enriched in the intermediate peak (Fig. 2C, S4-S6) (*9, 12, 21*). Consistent with these findings, some mutations at the phosphorylation sites or the Walker nucleotide binding motif in CII have been previously shown to create low amplitude or dampened oscillations (*22*). Here we show that these mutational effects are widespread (Fig. S5), suggesting that even when catalysis is compromised, KaiC can respond to KaiA and allosterically transmit output signals to create transcriptional rhythms. The KaiC protein sequence is highly conserved within cyanobacteria (*23*), and a natural question is how conservation correlates with phenotype. The phenotypes of substitutions at highly conserved positions are highly enriched for loss of amplitude, suggesting that selection is acting to maintain high amplitude self-sustained rhythms (Fig. 2D).

Mutating the interaction surfaces involved in the post-translational feedback loop controlling KaiC phosphorylation can cause loss of amplitude, in particular the groove in the KaiA C-terminal domain where KaiC binds for activation during the day (*9, 24*) (Fig. 2E) and the KaiA surface where KaiB binds for inactivation during the night (*12*) (Fig. 2F, S4-S6). These interactions are needed to sustain the in vitro phosphorylation oscillation, again indicating the importance of this post-translational system for high amplitude rhythms in vivo.

### Segregation of long-period and short-period phenotypes across genes

A fundamental question is how the 24-hour period is encoded in the structure of clock proteins. We find many mutations that change period while retaining high amplitude. Remarkably, the effect of mutation on period segregates sharply between the Kai proteins. Mutations in KaiA almost exclusively cause period lengthening, and mutations in KaiB are strongly enriched for period shortening. Mutations in KaiC can cause both effects, as well as rare period mutations with very large effect size (Fig. 3A-B).

**Figure 3.**
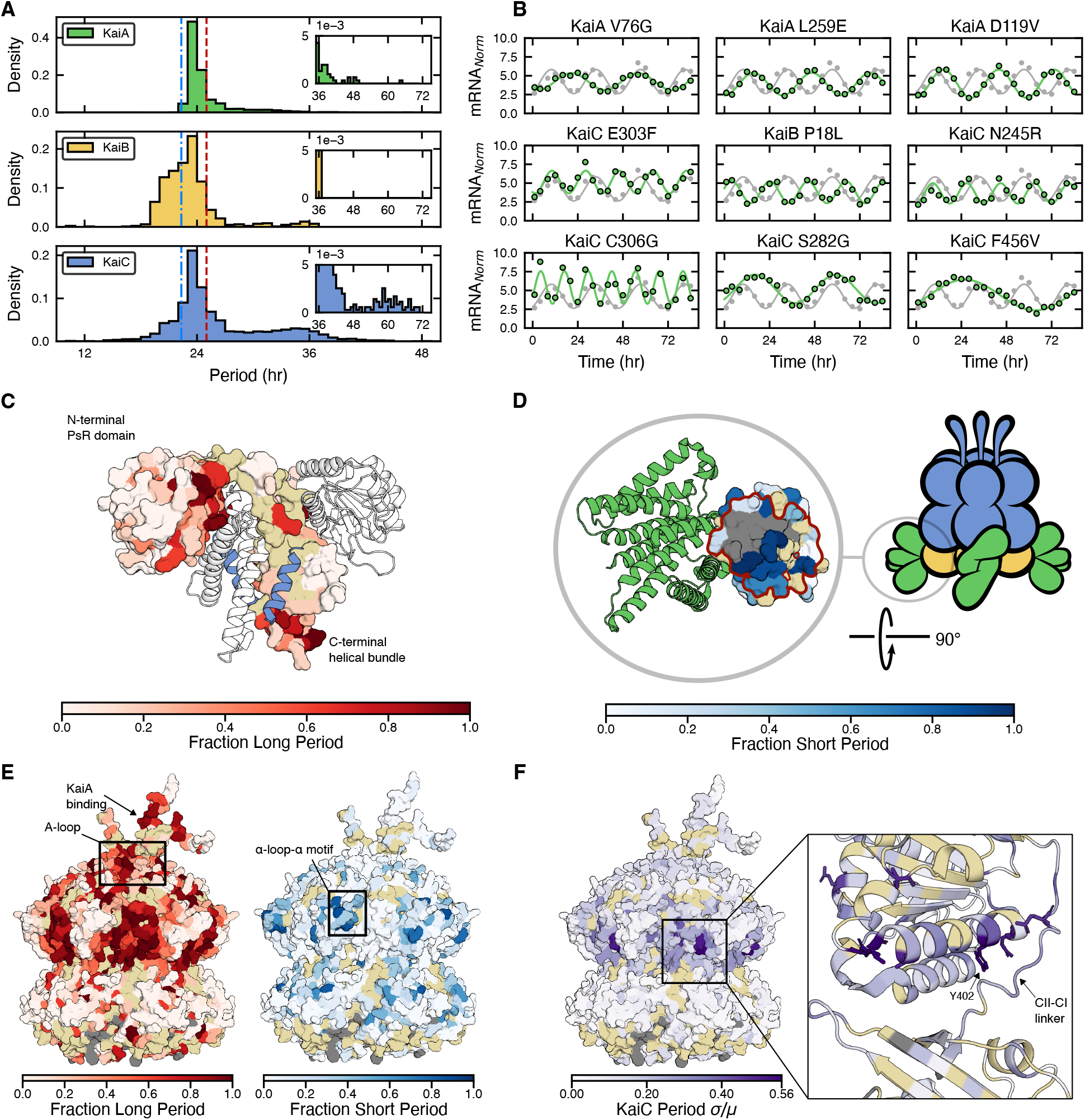
Localization of period mutations in the Kai protein structures. (**A**) Distribution of mutant periods by protein. Insets show the long period tails of the histograms. Dashed vertical lines represent 4 standard deviations above (*red*) or below (*blue*) the mean of subsampled WT used to define short period (< 22.33 hr) and long period (> 25.01 hr). (KaiA: n=3,046, KaiB: n=730, KaiC: n=3,580 (see Fig. S2). (**B**) Example traces of selected period mutants. Top row: long period mutants in KaiA (KaiA V76G = 35.5 hr, KaiA L259E = 31.0 hr, KaiA D119V = 29.5 hr). Middle row: short period mutants (KaiC E303F = 17.9 hr, KaiB P18L = 20.3 hr, KaiC N245R = 18.1 hr). Bottom row: extreme period mutations in KaiC (KaiC C306G = 12.6 hr, KaiC S282G = 43.0 hr, KaiC F456V = 74.6 hr) (**C-E**) Protein structures (*12, 21, 47, 48*) colored by fraction of high amplitude mutations belonging to short or long period groups. Beige: fewer than 5 mutants have a defined period. Gray: fewer than 5 mutants present. (**C**) One subunit of KaiA dimer (5C5E) colored by fraction of mutations causing long period. The other subunit is shown in semi-transparent ribbon representation. Blue: C-terminal fragment of KaiC bound to KaiA (**D**) The surface of KaiB in the nighttime KaiABC complex (5JWR) colored by fraction of mutations causing short period. View is from the KaiC CI domain looking down onto KaiB. Green ribbon: KaiA C-terminal domain. Area within the red outline: interface with KaiC (**E**) KaiC hexamer structure (3DVL) colored by fraction of long period mutants (*left*) and fraction of short period mutants (*right*). Annotations mark the A-loop, exposed by KaiA binding (*49*) and an α-loop-α motif implicated in nucleotide exchange (*28*) (**F**) KaiC hexamer structure colored by the standard deviation divided by the mean of the distribution of periods at each position. Licorice representation of side chains was added to residues that have relative standard deviation larger than 0.4. We call out Y402, a previously characterized position with large period tunability (*18*).

Long period mutations in KaiA are found surrounding the binding site for the KaiC C-terminal tail, as well as concentrated in the *pseudo*-receiver domain near the hinge in the KaiA dimer (Fig. 3C). Since this hinge rotates markedly in the inhibitory KaiABC complex, mutations here may affect the interconversion between the active and inhibited forms of KaiA (*12*). Notably, period lengthening of a similar magnitude can be caused in the reconstituted oscillator by reducing the KaiA dosage in the reaction (*25, 26*), suggesting that KaiA period mutations may be interpreted as reducing the effective KaiA activity.

Short period mutations in KaiB are highly enriched for the binding surface where the KaiB•KaiC complex forms (Fig. 3D, S7-8). The assembly and disassembly of this complex is known to be slow and coupled to the ATPase reaction in the KaiC CI domain (*10, 27*). Thus, our results suggest an important kinetic role for the KaiB•KaiC complex to prevent premature dissociation and maintain oscillator period.

Mapping period mutations to the KaiC structure confirms that residues near the KaiA binding site, and the A-loop whose exposure is regulated by KaiA binding, can cause period lengthening (Fig. 3E, S7-8). However, this analysis also reveals a striking hot spot of long period mutations on the KaiC CII domain near the waist of the double-domain hexamer. In addition, an α-loop-α motif on the exterior of CII near the catalytic site is enriched for short period mutations. This motif was previously implicated in the nucleotide exchange process (*28*).

### Interdomain regulation, not CI active site mutations, drives most period variation

To determine which positions in the Kai proteins have the strongest effect on period, we scored positions by the relative standard deviation of the distribution of oscillator periods caused by each amino acid substitution. This analysis reveals a region proximal to the CI-CII linker, including the α8 helix, that potently tunes oscillator period when mutated, causing a broad distribution of both shortening and lengthening (Fig. 3F). This analysis also implicates residues between the nucleotide binding site in CII and the central channel of the hexamer (Fig. S9). Similar properties were previously reported for mutation of Tyr402 in KaiC (*18*), which we now reveal to be representative of a localized group of positions with strong period tuning effects (Fig. S10). Previous work has shown that many period mutations in KaiC proportionally change the ATP turnover rate in the CI domain (*19, 29*). However, our global view of period mutations indicates that the most effective way to change period mutationally is to act distal to the CI domain presumably by altering interdomain regulation.

### Period can be tuned without compromising waveform or amplitude

By observing thousands of mutants in parallel, we are able to define a correlation structure for the dynamical properties of the oscillator phenotype. Mathematical models of oscillator topologies are known produce distinct amplitude-period correlations when randomly perturbed, depending on e.g. the presence of positive feedback loops (*30*). Empirically, we find that high amplitude rhythms are readily produced in both short and long period mutants (Fig. S12). Further, we find that mutant oscillators in KaiABC generally retain a nearly balanced waveform, in contrast to models of relaxation oscillators where parameter variation can produce severe asymmetry between rising and falling phases (*31*) (Fig. S11). Thus, the period of oscillation in the Kai system can be tuned by mutation without compromising amplitude or waveform shape.

However, mutations that cause partial loss of amplitude while retaining clearly detectable rhythms often access a specific non-circadian mode with long period rhythms centered around ∼34 hours (Fig. S12). Mutants with this property are enriched around the catalytic site in the KaiC CII domain, compared to the CI domain where mutations tend to destroy regular rhythmicity (Figs. S13, S14). This suggests an underlying weak but defined rhythmicity in the system that can operate when KaiC phosphorylation is not functional (*22, 32*).

### Low dimensional phase-period relationship suggests an entrainment boundary

The oscillating output of a circadian clock is characterized by a phase angle which indicates when peak output occurs relative to the light-dark cycles to which it is entrained. This quantity is likely physiologically very important because it determines the match between the gene expression program and the external environment. Plotting phase against period reveals a low dimensional structure in the high amplitude phenotypes accessible by single mutations (Fig. 4A). Many mutants can be found in a central peak that is not statistically distinguishable from wildtype. Away from this peak, period mutants follow a defined quasi-1D arc in the plane where phase increases sharply as period falls below ∼22 hours and apparently terminates near 18 hours. In contrast, mutants that create low amplitude oscillations lack a smoothly tunable period and cluster in the phase-period plane near a ∼34 hour period (Fig. 4B).

**Figure 4.**
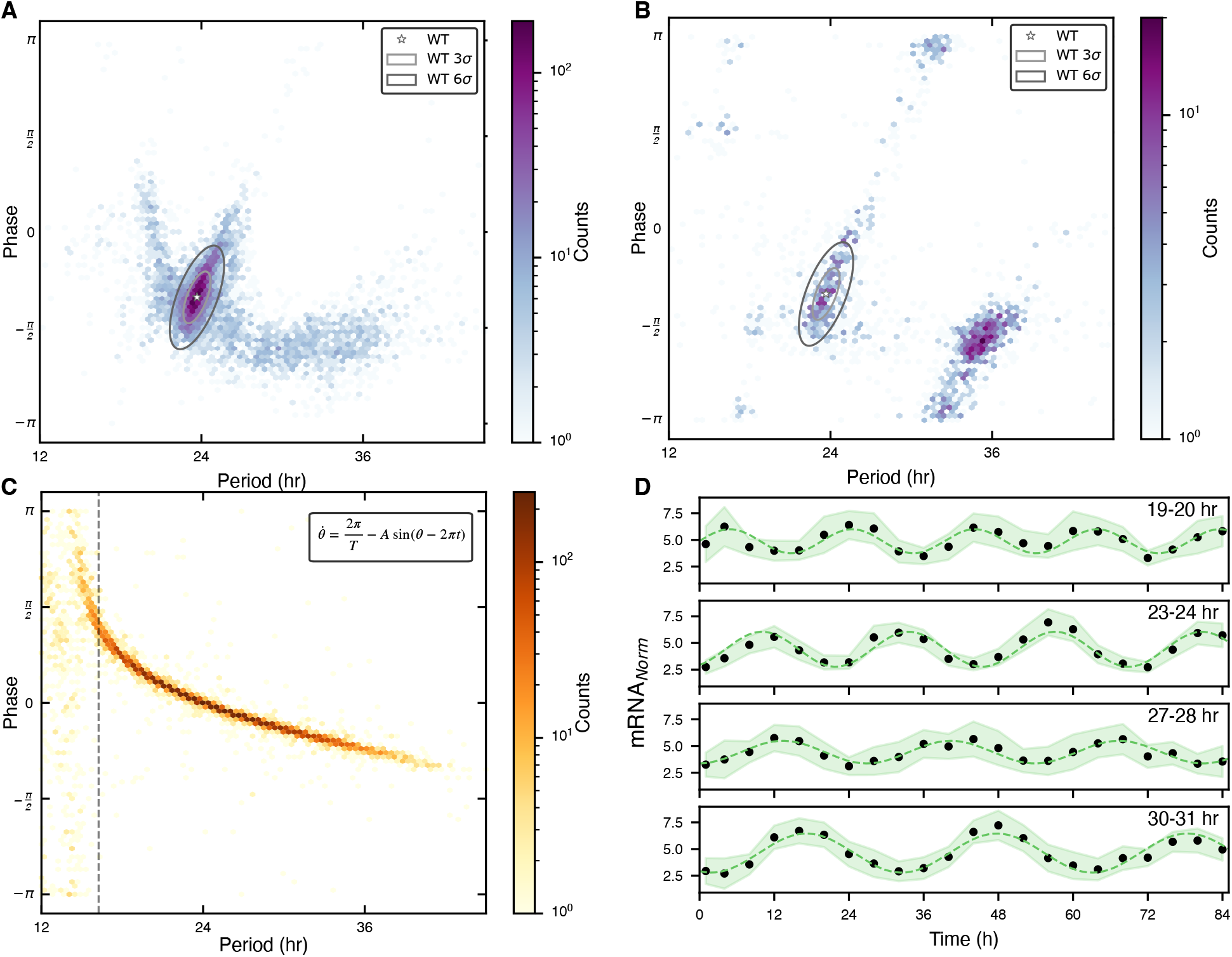
Low dimensionality of the phenotypic landscape. (**A**) Phase vs period for high amplitude mutants with defined period (n=7,356). Ellipses show 3-sigma and 6-sigma boundaries on fit parameters for subsampled WT barcodes. (**B**) Phase vs period for low amplitude mutants with defined period (n=2,089). Ellipses as in panel (A). (**C**) Phase-period relationship in a minimal phase oscillator model with Kuramoto coupling. Mutations are simulated by drawing oscillator periods (*T*) and coupling constants (*A*) from Gaussian distributions with 10x more variability in period (mean coupling constant A = 0.125 / day). Dashed line marks the loss of phase-locking (T = 16.2 hours) in model with mean parameters. (A-C) All color bars are logarithmic. (**D**) Average waveforms for high amplitude mutants with periods in the specified range. Dashed line shows sinusoidal model fit. Shading indicates standard deviation in the data. Note systemic variation of phase with period.

The regular pattern of phase and waveform in the high amplitude mutants can be visualized by averaging data from mutant oscillators with similar period together (Fig. 4D).We interpret these results using a minimal mathematical model of a driven stable oscillator, where a phase angle describes an oscillator that moves with constant internal velocity and Kuramoto coupling to an external environment (*33, 34*). This model has two parameters: a free-running period *T* and a coupling constant *A*. Though the fine structure of the data is not captured by this model, when we simulate mutation by putting 90% of the variability into the period (*T)*, we observe a qualitatively similar arc that increases in phase with shortened period until terminating at the boundary where phase-locking is lost (Fig. 4C). Thus, from the landscape formed by our Kai mutant library we predict that most mutations that change the period of the clock do so without substantially altering the sensitivity to external light-dark cycles.

## Discussion

The statistical structure of phenotypes accessible by mutation may indicate the history of the evolutionary demands on the system (*35*) and can put constraints on the underlying molecular mechanisms. The overall picture that emerges for this circadian clock is that of a system with remarkable but defined mutational plasticity. The structural localization of phenotypes suggests that many effects can be understood as altered interactions between and within the Kai proteins, though, because we are studying this system in its native context, some phenotypes may involve altered interaction with other clock components. (*12, 36, 37*)

Only a minority of mutations (31%) are neutral, producing both amplitude and period similar to wildtype. Many mutations at highly conserved positions cause either arrythmia or a severe loss of amplitude, indicating that sustained oscillation is a fragile system property in certain sequence dimensions that requires multiple elements, including the KaiA-dependent feedback loop, to function. However, along other dimensions of variation, period emerges as a surprisingly mutable direction in phenotypic space. Many mutations apparently change free-running period without noticeably impacting amplitude, waveform, or coupling to the external environment. This result is unexpected for two reasons: first because many possible oscillator architectures do not have this property, and second because the period of the day-night cycle is near-invariant over deep evolutionary time.

The ability of biological systems to maintain essential functions under constant selection pressures while retaining the capacity for phenotypic innovation has been noted at scales from atoms (*38-40*) to ecosystems (*41, 42*). The central design principle that encodes this characteristic is also becoming clear—a sparse, conserved, mutationally-sensitive core of strong interactions takes care of essential functionality, enabling a mutationally-tolerant, fungible environment to deliver smooth phenotypic change (*40, 43*). This architecture enables cryptic phenotypic variation to accumulate in populations even without selection, a property that has been argued to promote the evolvability of biological systems (*42, 44, 45*). The data presented here are consistent with this principle—the Kai system displays a conserved core essential for oscillation and a mutable surround enabling many paths to smoothly traverse the arc of phase-period covariation while maintaining high amplitude oscillation. This tunability of the extant system may reflect the evolutionary path used to precisely set the catalytic rate of KaiC from a RecA/DnaB superfamily ancestor with a non-circadian function (*46*). In contrast, the mutations that shift to low amplitude oscillations lack tunability and cluster in a long period mode.

These architectural principles have implications for making physical models for biological systems. Many plausible mechanisms may account for functional characteristics, but it is possible that only some will be consistent with the structure of the phenotypic landscape exposed by studies such as this one. We propose that the statistics of variation under mutation, enabled by high-throughput DNA synthesis and sequencing, may provide key constraints for making unified models for biological systems — ones that simultaneously account for physiological function, biophysical mechanism, and evolvability.

## Acknowledgments

We thank members of the Rust and Ranganathan labs for advice and discussion. We thank Jenny Lin for preliminary development of the method and Gopal Pattanayak for design of plasmids.

## Author contributions

Conceptualization: DS, RR, MJR

Methodology: SJK, DS, BA, RR, MJR

Investigation: SJK, DS, BA, RR, MJR

Funding acquisition: RR, MJR

Project administration: RR, MJR

Supervision: RR, MJR

Writing – original draft: SJK, DS, MJR

Writing – review & editing: SJK, DS, BA, RR, MJR

## Data, code, and materials availability

All data are available in the main text or the supplementary materials. Custom sequencing analysis and data analysis software available on request.

## Supplementary Materials

Materials and Methods Supplementary Text

Figs. S1 to S14

Table S1 to S2

Data S1 to S2

## References and Notes

1. M. A. Stiffler, D. R. Hekstra, R. Ranganathan, Evolvability as a function of purifying selection in TEM-1 beta-lactamase. Cell 160, 882–892 (2015).

2. P. Bandaru et al., Deconstruction of the Ras switching cycle through saturation mutagenesis. Elife 6, (2017).

3. M. Ishiura et al., Expression of a gene cluster kaiABC as a circadian feedback process in cyanobacteria. Science 281, 1519–1523 (1998).

4. M. Nakajima et al., Reconstitution of circadian oscillation of cyanobacterial KaiC phosphorylation in vitro. science 308, 414–415 (2005).

5. J. S. Markson, J. R. Piechura, A. M. Puszynska, E. K. O’Shea, Circadian control of global gene expression by the cyanobacterial master regulator RpaA. Cell 155, 1396–1408 (2013).

6. M. J. Rust, J. S. Markson, W. S. Lane, D. S. Fisher, E. K. O’Shea, Ordered phosphorylation governs oscillation of a three-protein circadian clock. Science 318, 809–812 (2007).

7. T. Nishiwaki et al., A sequential program of dual phosphorylation of KaiC as a basis for circadian rhythm in cyanobacteria. EMBO J 26, 4029–4037 (2007).

8. H. Iwasaki, T. Nishiwaki, Y. Kitayama, M. Nakajima, T. Kondo, KaiA-stimulated KaiC phosphorylation in circadian timing loops in cyanobacteria. Proc Natl Acad Sci U S A 99, 15788–15793 (2002).

9. I. Vakonakis, A. C. LiWang, Structure of the C-terminal domain of the clock protein KaiA in complex with a KaiC-derived peptide: implications for KaiC regulation. Proc Natl Acad Sci U S A 101, 10925–10930 (2004).

10. C. Phong, J. S. Markson, C. M. Wilhoite, M. J. Rust, Robust and tunable circadian rhythms from differentially sensitive catalytic domains. Proc Natl Acad Sci U S A 110, 1124–1129 (2013).

11. Y. G. Chang, R. Tseng, N. W. Kuo, A. LiWang, Rhythmic ring-ring stacking drives the circadian oscillator clockwise. Proc Natl Acad Sci U S A 109, 16847–16851 (2012).

12. R. Tseng et al., Structural basis of the day-night transition in a bacterial circadian clock. Science 355, 1174–1180 (2017).

13. H. Kageyama et al., Cyanobacterial circadian pacemaker: Kai protein complex dynamics in the KaiC phosphorylation cycle in vitro. Mol Cell 23, 161–171 (2006).

14. A. Gutu, E. K. O’Shea, Two antagonistic clock-regulated histidine kinases time the activation of circadian gene expression. Mol Cell 50, 288–294 (2013).

15. I. Mihalcescu, W. Hsing, S. Leibler, Resilient circadian oscillator revealed in individual cyanobacteria. Nature 430, 81–85 (2004).

16. J. Chew, E. Leypunskiy, J. Lin, A. Murugan, M. J. Rust, High protein copy number is required to suppress stochasticity in the cyanobacterial circadian clock. Nat Commun 9, 3004 (2018).

17. E. M. Clerico, J. L. Ditty, S. S. Golden, Specialized techniques for site-directed mutagenesis in cyanobacteria. Methods Mol Biol 362, 155–171 (2007).

18. K. Ito-Miwa, Y. Furuike, S. Akiyama, T. Kondo, Tuning the circadian period of cyanobacteria up to 6.6 days by the single amino acid substitutions in KaiC. Proceedings of the National Academy of Sciences 117, 20926–20931 (2020).

19. J. Abe et al., Circadian rhythms. Atomic-scale origins of slowness in the cyanobacterial circadian clock. Science 349, 312–316 (2015).

20. H. Nishimura et al., Mutations in KaiA, a clock protein, extend the period of circadian rhythm in the cyanobacterium Synechococcus elongatus PCC 7942. Microbiology (Reading) 148, 2903–2909 (2002).

21. R. Pattanayek et al., Visualizing a circadian clock protein: crystal structure of KaiC and functional insights. Mol Cell 15, 375–388 (2004).

22. Y. Kitayama, T. Nishiwaki, K. Terauchi, T. Kondo, Dual KaiC-based oscillations constitute the circadian system of cyanobacteria. Genes Dev 22, 1513–1521 (2008).

23. V. Dvornyk, O. Vinogradova, E. Nevo, Origin and evolution of circadian clock genes in prokaryotes. Proc Natl Acad Sci U S A 100, 2495–2500 (2003).

24. R. Pattanayek et al., Analysis of KaiA-KaiC protein interactions in the cyano-bacterial circadian clock using hybrid structural methods. EMBO J 25, 2017–2028 (2006).

25. A. G. Chavan et al., Reconstitution of an intact clock reveals mechanisms of circadian timekeeping. Science 374, eabd4453 (2021).

26. M. Nakajima, H. Ito, T. Kondo, In vitro regulation of circadian phosphorylation rhythm of cyanobacterial clock protein KaiC by KaiA and KaiB. FEBS Lett 584, 898–902 (2010).

27. Y. G. Chang et al., Circadian rhythms. A protein fold switch joins the circadian oscillator to clock output in cyanobacteria. Science 349, 324–328 (2015).

28. L. Hong, B. P. Vani, E. H. Thiede, M. J. Rust, A. R. Dinner, Molecular dynamics simulations of nucleotide release from the circadian clock protein KaiC reveal atomic-resolution functional insights. Proc Natl Acad Sci U S A 115, E11475–E11484 (2018).

29. K. Terauchi et al., ATPase activity of KaiC determines the basic timing for circadian clock of cyanobacteria. Proc Natl Acad Sci U S A 104, 16377–16381 (2007).

30. T. Y. Tsai et al., Robust, tunable biological oscillations from interlinked positive and negative feedback loops. Science 321, 126–129 (2008).

31. R. FitzHugh, Impulses and physiological states in theoretical models of nerve membrane. Biophysical Journal 1, 445–466 (1961).

32. X. Qin, M. Byrne, Y. Xu, T. Mori, C. H. Johnson, Coupling of a core post-translational pacemaker to a slave transcription/translation feedback loop in a circadian system. PLoS Biol 8, e1000394 (2010).

33. G. B. Ermentrout, J. Rinzel, Beyond a pacemaker’s entrainment limit: phase walk-through. Am J Physiol 246, R102–106 (1984).

34. S. Strogatz, Nonlinear dynamics and chaos: with applications to physics, biology, chemistry, and engineering. (CRC Press, Boca Raton, ed. Third edition., 2024), pp. pages cm.

35. N. Kashtan, U. Alon, Spontaneous evolution of modularity and network motifs. Proc Natl Acad Sci U S A 102, 13773–13778 (2005).

36. R. Tseng et al., Cooperative KaiA-KaiB-KaiC interactions affect KaiB/SasA competition in the circadian clock of cyanobacteria. J Mol Biol 426, 389–402 (2014).

37. S. J. Kim, C. Chi, G. Pattanayak, A. R. Dinner, M. J. Rust, KidA, a multi-PAS domain protein, tunes the period of the cyanobacterial circadian oscillator. Proc Natl Acad Sci U S A 119, e2202426119 (2022).

38. S. Bershtein, M. Segal, R. Bekerman, N. Tokuriki, D. S. Tawfik, Robustness-epistasis link shapes the fitness landscape of a randomly drifting protein. Nature 444, 929–932 (2006).

39. E. J. Hayden, E. Ferrada, A. Wagner, Cryptic genetic variation promotes rapid evolutionary adaptation in an RNA enzyme. Nature 474, 92–95 (2011).

40. A. S. Raman, K. I. White, R. Ranganathan, Origins of Allostery and Evolvability in Proteins: A Case Study. Cell 166, 468–480 (2016).

41. S. E. Luria, M. Delbruck, Mutations of Bacteria from Virus Sensitivity to Virus Resistance. Genetics 28, 491–511 (1943).

42. A. Wagner, Neutralism and selectionism: a network-based reconciliation. Nat Rev Genet 9, 965–974 (2008).

43. M. Kirschner, J. Gerhart, Evolvability. Proc Natl Acad Sci U S A 95, 8420–8427 (1998).

44. J. A. Draghi, T. L. Parsons, G. P. Wagner, J. B. Plotkin, Mutational robustness can facilitate adaptation. Nature 463, 353–355 (2010).

45. J. A. Draghi, J. B. Plotkin, Molecular evolution: Hidden diversity sparks adaptation. Nature 474, 45–46 (2011).

46. A. Mukaiyama et al., Evolutionary origins of self-sustained Kai protein circadian oscillators in cyanobacteria. Nat Commun 16, 4541 (2025).

47. S. Ye, I. Vakonakis, T. R. Ioerger, A. C. LiWang, J. C. Sacchettini, Crystal structure of circadian clock protein KaiA from Synechococcus elongatus. J Biol Chem 279, 20511–20518 (2004).

48. R. Pattanayek, M. Egli, Protein-Protein Interactions in the Cyanobacterial Circadian Clock: Structure of KaiA Dimer in Complex with C-Terminal KaiC Peptides at 2.8 A Resolution. Biochemistry 54, 4575–4578 (2015).

49. Y. I. Kim, G. Dong, C. W. Carruthers, Jr., S. S. Golden, A. LiWang, The day/night switch in KaiC, a central oscillator component of the circadian clock of cyanobacteria. Proc Natl Acad Sci U S A 105, 12825–12830 (2008).

