## Supplemental Information for "The Phenotypic Landscape of a Circadian Clock"

### Materials and Methods

#### Construction of the *kaiABC* DMS plasmid library:

The coding sequences of *kaiABC* were divided into 24 total subsequences, hereafter “tiles,” of exactly 120 base pairs. For each tile, we designed 40 oligonucleotides, one for each codon to be mutated, except for the last tile in each gene, which had fewer codons and was buffered by unmutated 3’ UTR sequence to the correct length. Each oligonucleotide had the tile coding sequence with one codon replaced by “NNK,” flanked on each side by ~20 base pairs of non-tile coding sequence, flanked on each side by common cloning adapters, flanked on each side by orthogonal primer binding sites (*1*). The 5’ and 3’ cloning adapters were CACCTGCAAGGTTCA and GGAACATGGCAGGTG, respectively.
For the synthesized oligo nucleotides, several synonymous edits were made to the *kaiABC* coding sequence to minimize synthesis inefficiencies. The edits are as follows, numbered according to the codon of each gene: *kaiA*(144:GGG->GGT), *kaiC*(71:GGG->GGT), *kaiC*(175:GGG->GGA), *kaiC*(213:GGG->GGT), *kaiC*(233:GGG->GGT), *kaiC*(250:GGG->GGA), *kaiC*(385:CGG->CGA), *kaiC*(477:CCG->CCC). Tiles were synthesized as oPool libraries at 1 pmol scale from IDT.

For each tile, the pool of oligonucleotides was PCR amplified using the orthogonal primers specific to that tile. The amplicons were then cloned into a pUC19-derived storage vector using PaqCI-based Golden Gate Assembly. For each tile, a pUC19-derived destination vector where AmpR is replaced with KanR was prepared by Gibson Assembly that contained the full *kaiABC* sequence with the corresponding tile deleted and replaced with a BspQI-flanked insert that encoded trimethoprim resistance. BspQI-based Golden Gate Assembly was used to scarlessly shuttle the mutated tile libraries from the storage vector to the destination vector, simultaneously removing the trimethoprim resistance marker. All 24 tile-harboring destination vectors were pooled at equal molarity and used as a template to PCR amplify the full *kaiABC* sequence with primers that introduced a synthetic sequence following the *kaiC* STOP codon with a 15 random bp barcode and also added new BspQI sites:

…**TAG***GCGATTAAGTTGGGTAACGCCAGGG*GTCTCAGCTCNNNNNNNNNNNNNNNGC
*GGCCGCCGATCCTCTAGTATGCTTG*

The KaiC STOP codon is in bold and the priming sites for sequencing are italicized.

The resulting amplicon was cloned into the neutral site targeting using BspQI-based Golden Gate Assembly along with 457 bp upstream of the *kaiA* coding sequence. The neutral site targeting vector is a modified pAM1303 (Addgene plasmid #40243) that has an additional AmpR gene added outside of the neutral site homology region for selection in *E. coli*, and CmR gene that is removed when the Golden Gate Assembly occurs. pAM1303 targets Neutral Site I in the *Synechococcus elongatus* genome and integrates a spectinomycin/streptomycin resistance cassette along with the mutated *kaiABC* gene cluster. At all steps, the pool size was maintained at >1000X the theoretical library size, except for the post-barcoding cloning, which was estimated at 40 barcodes per variant.

The plasmid library was linearized by blunt end digestion with ScaI and cleaned up using a DNA purification kit (Zymo DNA Clean and Concentrator). The linearized plasmid preparation was used for commercial long read sequencing based on rolling-circle amplification (SeqCenter, PacBio Revio flow cell).

To associate barcodes with mutations, raw reads were trimmed to the kaiABC coding sequences plus the barcode, and reads in the wrong orientation were transformed to their reverse-complement.710,779 barcodes were mapped using an error-correcting barcode mapping algorithm (BCAR) (*2*). Briefly, the algorithm performs a multiple sequence alignment for all reads with the same barcode. For each group of aligned reads, a consensus read is generated. For each position, a quality score is estimated by comparing the raw base calls and quality scores at each aligned position. The consensus reads were then filtered to only keep reads with a minimum quality score >30 and with more than 2 reads comprising that consensus sequence. Finally, mutations in the *kaiABC* sequence were called by comparing the consensus alignment for each barcode to the wildtype reference sequence.

### Transformation of *S. elongatus* with the DMS plasmid library

1L of an *S. elongatus* strain where the *kaiABC* cluster had been knocked out with a gentamycin resistance casette was grown in 4 acid washed 1L flasks with 250mLs of BG11+20mM HEPES pH 8 to a target OD750 of 0.5 at ~50 µmol photons $m^{-2}s^{-1}$ from white LEDs. The culture was then spun down in 500mL centrifuge tubes at 5020xg for 10 min. Each pellet was resuspended in 500mL of 10 mM NaCl then spun down again. This washed pellet was then resuspended in 80mL BG11 to a target OD750 of 5.0. 40 µg of a KaiABC plasmid library maxiprep (Qiagen endofree plasmid maxi kit) was then combined with the culture in a 250mL acid-washed Erlenmeyer flask. The flask was wrapped in foil and incubated with shaking at 30 C 190 RPM overnight.

The next day the culture was resuspended in 15mL BG11. A dilution series was plated on selective BG11 1.4 % gelzan plates to estimate transformation efficiency. The remaining recovery was plated on 15 245 mm Square BioAssay dishes (Corning, 431111) containing BG11 1.4 % gelzan 2ug/mL spectinomycin/streptomycin solid culture medium, aliquoting 1 mL of the transformation mixture per plate and spreading with glass beads. Plates were wrapped with vent tape (3M 394) and grown at ~50 µmol photons $m^{-2}s^{-1}$ from LEDs. Once visible colonies had appeared (about 7 days), we estimated the number of transformants to be ~4 million. 10mL of BG11 sp/st. was added to each plate and an L spreader was used to resuspend colonies into the liquid. To test for contamination with heterotrophic microbes, resuspensions were spotted onto standard LB-agar plates and incubated at 30 C for 2 days. Resuspensions whose LB test plates showed no growth were pooled as the final library. The cultures for time course sampling were inoculated directly from this pool.

### Time-course sampling and nucleic acid extraction:

The cyanobacterial DMS library was inoculated in BG11 to a total volume of 900 mL at OD750 of 0.02. The culture was grown at 30°C with shaking at 185 RPM in a Percival incubator under constant illumination of ~100 µmol photons $m^{-2}$ $s^{-1}$ from cool white fluorescent bulbs, divided into 9 side-by-side flasks each containing 100 mL of culture. When the OD750 reached ~ 0.3, the cultures were subjected to two cycles of 12-h light and 12-h dark for entrainment. After the 2-day entrainment, the cultures were released back into constant light (t = 0). Cells were sampled every 4 hours in constant light (t = 4 h, 8 h, …, 84 h), except for the first time point, which was taken at t = 1 h. 90 mL of culture was sampled at each time point by drawing 10 mL from each flask, which were pooled then harvested by vacuum filtration onto cellulose nitrate filter papers with 0.45 µm pore size (Whatman #10401170). The filter paper was placed into a 50 mL conical tube, flash frozen in liquid nitrogen, and stored at –80°C. The cultures were manually held at OD750 ~ 0.3 by back dilution. The average doubling time during sampling in constant light from was 22 ± 4.5 hours.  Once per day, at 3 PM, an additional 9 mL of culture was collected by drawing 1 mL from each flask for DNA sequencing and pelleted by centrifugation, followed by liquid nitrogen freezing and storage at –80°C.

RNA extraction was conducted using RNase-free barrier pipette tips, tubes, and reagents. Gloves, pipettes, microcentrifuges, and bench surface were wiped with RNase decontamination solution (RNAse AWAY, Thermo Scientific).  Samples were processed in batches of 12 or fewer to minimize sample-to-sample differences in handling delays. 50 mL conical tubes containing the cell pellets were removed from the –80°C freezer and placed on ice, the cell pellet touching wall facing down. Using a 1000 µL pipette, 1 mL of room temperature Trizol (Invitrogen, 15596026) was ejected directly onto the cell pellet on the filter paper, followed by multiple rinsing of the filter paper by drawing the Trizol that collects at the bottom of the conical tube. Resuspended cells were placed on ice for the Trizol resuspension to collect at the bottom, while other samples are handled in the same way. Each sample was then transferred to a 2 mL screw-cap tube pre-filled with 0.1 mm acid-washed glass beads (Benchmark Scientific, D1031-01), which were vortexed at room temperature at 3000 RPM for cell lysis. After vortexing, 200 µL of chloroform was added to each tube, which was vigorously mixed by hand for 15 seconds and then incubated at room temperature for 2–3 minutes. Samples were then centrifuged at 12,000 g for 15 minutes at 4°C. From each sample, the upper aqueous phase was carefully decanted without disturbing the interphase and transferred to a clean microcentrifuge tube. The volume of the recovered aqueous phase was noted and 1.25x volumes of ice-cold isopropanol was added to the tube to precipitate RNA. After incubation at room temperature for 10 minutes, samples were centrifuged at 15,000 g for 10 minutes at 4°C. Without disturbing the RNA pellet, the supernatant was decanted by pipette and discarded. The pellet was resuspended by adding 700 µL 75% ice-cold ethanol to the tube and mixed by pipetting up and down, ensuring that precipitated RNA on the wall of the tube is also being resuspended, followed by vortexing. Samples were then centrifuged at 7500 g for 5 minutes at 4°C. Without disturbing the RNA pellet, supernatant in each tube was carefully removed in two steps, first using a 1000 µL pipette and then using a 200 µL pipette. RNA pellets were then air-dried until the supernatant was mostly evaporated but while the pellet still retains a wet appearance. Each pellet was resuspended in 30 µL RNase-free water by pipetting up and down, followed by incubation at 60°C incubation for 7 minutes. After resuspension, RNA sample containing tubes were placed on ice. Nucleic acid concentrations were measured by absorbance at 260 nm using a Nanodrop spectrophotometer (Thermo Scientific). Samples were stored at –80°C.

To process frozen cells for DNA extraction, cell pellets were resuspended in 200 µL 10mM Tris pH 8, and added to screw cap Eppendorf tubes with 100um acid-washed beads added. 250 µL of 25:24:1 phenol:chloroform:IAA was added, and the tubes were vortexed for 5 minutes at max speed. Samples were then spun down for 10 minutes at max speed in a microcentrifuge at room temperature. The aqueous phase was then carefully decanted and and transferred to a clean tube. DNA precipitation was performed by adding 22.2 µL of 3 M NaOAc and 450 µL ethanol, mixing by inversion, and centrifugation for 15 minutes at 20,000 g at 4C. The supernatant was removed and the pellet was washed twice with 1 mL 70% ethanol. After a final spin for 5 minutes at 20,000 g at 4C, the supernatant was removed and the pellet was allowed to dry. The pellet was then resuspended in 30uL 10 mM Tris pH 8.

**Reverse transcription and sequencing:**

After thawing on ice, 10 µg of the RNA sample (or the entirety of the extracted RNA for less concentrated samples) was treated with Turbo DNase (Invitrogen AM1907) to remove contaminating genomic DNA. The DNase reaction was prepared by mixing the RNA sample, 10x Turbo DNase buffer, and 3 µL Turbo DNase in a 35 µL reaction. The DNase was added in two steps. 1.5 µL was added first, followed by 30 minutes of 37°C incubation, at which point 1.5 µL more was added. The sample was then incubated at 37°C for 30 more minutes. The reactions were treated with 7 µL of inactivation slurry by room temperature incubation for 2–5 minutes during which the samples were gently vortexed for resuspension. The samples were then centrifuged at 20,000 g for 2 minutes. 33 µL of supernatant was recovered and transferred to new microcentrifuge tubes. The concentrations of the RNA samples were measured by absorbance at 260 nm using a Nanodrop spectrophotometer. Samples were stored at –80°C.

DNase-treated RNA samples were reverse-transcribed using SuperScript IV reverse transcriptase (SSIV) (Invitrogen 18090050). Up to 1.3 µg of extracted RNA was used for each reaction, where the entire sample was used for timepoints with low RNA yield. Where possible, the total reaction volume was set at 20 µL, but for less concentrated samples, the reaction was scaled up as needed. A 1x reaction was prepared with RNA input, 1 µL of 50 mM random hexamer primers (New England Biolabs S1230S), 1 µL dNTPs, 4 µL 5x SSIV buffer, 1 µL 100 mM DTT, 0.5 µL Murine RNase inhibitor (New England Biolabs M0314S), 1 µL SSIV, and 1 µL RNaseH. RNA, primers, and dNTPs were mixed together in PCR tubes and incubated at 65°C for 5 minutes, followed by incubation on ice for 1 minute. A master mix of buffer, DTT, RNase inhibitor, and the enzyme was prepared and added to the primer-bound samples. The full reverse transcription reaction was incubated at 23°C for 10 minutes, 55°C for 10 minutes, and 80°C for 10 minutes to denature the RNAse inhibitor, followed by RNaseH reaction at 37°C for 20 minutes. A negative control reaction was also prepared with the same components, except the reverse transcriptase was replaced with water. All RT– samples were prepared for the total volume of 20 µL using a smaller amount of RNA for lower yield samples. The resulting cDNA samples were stored at –20°C. Intense bands from RT+ samples and no band or very faint bands from the RT– sample were confirmed by agarose gel analysis of 25 µL, 25x test PCR reactions with 1 µL of the cDNA sample and 0.25 µL of Taq polymerase.

gDNA and cDNA samples were quantified by Qubit hsDNA and ssDNA kits respectively (Thermo Fisher). Each sample was amplified in 5 cycles of PCR with Q5 Hot Start High Fidelity polymerase (NEB, M0493). 100 µg of gDNA was used as template in a 50uL PCR. For cDNA a target of 1 µg template was used (or, if the yield of the reverse transcription reaction was below 1 µg , the entire reaction product was used). The PCR was scaled up by 50 µL increments for each reaction as needed so that the template cDNA was no more than 1/5 the final volume. If the PCR was scaled up it was split into 50 µL subvolumes on the PCR plate. Round one primers were:

ACTCTTTCCCTACACGACGCTCTTCCGATCT(Ns)GCGATTAAGTTGGGTAACGCCAG and:

ACTGGAGTTCAGACGTGTGCTCTTCCGATCT(Ns)CAAGCATACTAGAGGATCGGCGG. These primers add on the Truseq adaptor sequence, and also add 4-7Ns that serve as unique molecular identifiers (UMIs).

Round 1 PCRs were cleaned up using PCRclean DX beads (Aline Biosciences). Post clean-up products were resuspended in 40 µL of 10mM Tris pH8, and at this point the split subvolumes were combined by resuspension into the same 40 µL volume. A second round of PCR added the remainder of the Illumina adaptor sequences. 10 µL of cleaned up round 1 PCR was used as template in a 20 cycle 50 µL Q5 PCR reaction. Using custom Illumina Truseq compatible primers (IDT). After PCR, the entire product was run on a 2% agarose gel and the final product (221-227bp) was gel extracted. The gel extractions were quantified using a Qubit hsDNA kit (Thermo Scientific) and mixed together in equimolar ratios. This mix was then quantified and diluted to 10 nM for sequencing on a NovaSeqX sequencer using a paired end 100 kit (Illumina).

**Data processing and time course analysis:**

Paired end reads were merged using FLASH (*3*) then the adaptor regions were trimmed using Cutadapt (*4*). Only trimmed reads of the correct length were kept. Then we analyzed the 15N barcode and the 4-7N UMIs on either end of the read. To account for PCR jackpotting we counted number of unique barcode UMI sets. The using the barcode map created by BCAR we assigned counts to each barcode. All redundant barcodes were pooled into counts for each mutant. RNA mutant counts were normalized by dividing by the number of DNA counts of that mutant. We then normalize that by the RNA counts/DNA counts of the null normalization mutants (see below for definition of the null group).

$$mRNA_{\text{norm}} = \frac{RNA\text{counts}_{\text{mutant}}}{DNA\text{counts}_{\text{mutant}}} / \frac{RNA\text{counts}_{\text{nulls}}}{DNA\text{counts}_{\text{nulls}}}$$

We fit the normalized RNA timecourses to the following transformed sine wave function:

$$y = S\cos\left(\frac{2\pi t}{T}\right) + C\sin\left(\frac{2\pi t}{T}\right) + B$$

Where $C = A\cos(\phi)$, $S = A\sin(\phi)$, A is amplitude, $\phi$ is phase, and B is baseline. We take a Fast Fourier Transform on the time series (numpy.fft), and take the 5 highest points in the discrete power spectrum as initial conditions for least squares regression. Least squares regression was performed using the lmfit Python library, and the fit with the lowest value of $\chi^2$ was selected.

A subset of nonsense mutations in KaiC were picked to be the null normalizations set that all mutants were divided by. First we normalized the KaiC nonsense mutants by the first 250 amino acids in KaiC and fit them as above to estimate amplitude. We selected a seed null set using an initial maximum amplitude threshold of 0.12, then renormalized and iterated this procedure until we obtained a self-consistent null set where all the amplitude of all nonsense mutants were < 0.3.

To determine filters for accurate parameter estimation we created a subset of WT barcodes. We took all WT barcodes and randomly drew 1-1000 barcodes to form a subset. We then fit these subsets as described above, testing different thresholds for the minimum number of counts per timepoint, and a minimum number of timepoints required to cross this threshold. We found that enforcing a threshold of 200 counts per timepoint in 20 out of 22 timepoints allowed > 94% of the WT barcode subsamples to estimate amplitude and baseline with <50% error compared to the full WT fit. 13,128 single missense mutants passed these filters and their amplitude and baseline were estimated from the sinusoidal regression described above.

To estimate period and phase, we set filters on the estimated error in the period and amplitude estimate returned from the least squares regression (square root of the local covariance matrix). Cutoffs of std(phase) < .695 and std(amplitude) / mean(amplitude) < .35 were tuned by hand, looking at examples of mutant fits that passed and failed the filter at extreme parameter values (high amplitude, low amplitude, etc).

To estimate waveform skew, we used a triangle wave model where one cycle of the oscillation is described by two piecewise linear segments, with a peak that occurs at a fraction α of the period:

$$f_{\text{tri}}(\tau) = \begin{cases} B + A\left(\dfrac{\tau}{\alpha T} - \dfrac{1}{2}\right), & \text{if } \tau < \alpha T \\ B - A\left(\dfrac{\tau - \alpha T}{\tau(1-\alpha)} + \dfrac{1}{2}\right), & \text{if } \tau > \alpha T \end{cases}$$

Where $\tau = t + T\left(\frac{\phi}{2\pi} + \frac{1}{4}\right) \bmod T$. Here, $T$ is the oscillator period, $\phi$ is the phase, $A$ is the amplitude, and $B$ is the baseline. We used least-squares regression (scipy.optimize.curvefit(), Trust Region Reflective method in Python) to fit this model to time series data, using the best-fit amplitude, baseline, period, and phase from sinusoidal fitting (described above) as initial conditions. Estimated values of the skewness parameter α from this process are shown in Fig. S10.

To generate simulated sinusoidal data and sawtooth data as shown in Fig. S10B, we selected amplitude, period, and baseline at random from the sinusoidal fits to the single mutant data classified as high amplitude (described above). These parameters were then used to generate either sine waves or sawtooth triangle waves (using the above function $f_{tri}$(t)) sampled at the same time points acquired in the experimental data. For sawtooth waves, α was drawn from a uniform distribution on (0, 1). Gaussian noise with $\sigma = 0.05\,\mu$ was added to each simulated data point, and the resulting simulated data sets were analyzed using the process described above.

**Assignment of mutant phenotypes and structural analysis:**

Missense mutants with a sufficient read depth were analyzed. Their amplitude distribution was fit using Gaussian mixture model, as shown in Figure 2A. The amplitude cutoff was determined as the minimum between the peak representing the low amplitude group and the peak representing the high amplitude group, which was 1.146. For all positions in each protein that have 5 or more sufficiently read substitutions, the fraction of the number of high amplitude mutants (Amplitude > amplitude cutoff) to the number of total mutants in the residue was calculated and color-mapped onto the protein structures, as shown in Figure 2. Of 9,445 single missense mutants passed that had sufficient read depth, 7,356 belonged to the high amplitude group, and 2,089 were low/null amplitude mutants.

The baseline of all single missense mutants was classified as low (<3.34), high (>5.14), or WT-like (>3.34 and <5.14). These cutoffs were determined as the mean of the subsampled WT distribution ±3x the standard deviation. The fraction high baseline or low baseline was calculated by dividing the number of high baseline or low baseline mutants by the total number of missense mutants with enough reads at that position. These values were color-mapped onto the protein structures, as shown in Figure S3.

Mutants in the high amplitude group that also pass the period error filter were used for period phenotype analysis for Figure 3. The cutoffs for short period and long period phenotypes were determined as the mean of the subsampled WT distribution +/– 4x standard deviation, which were 22.33 hr and 25.01 hr, respectively. The fraction short period (SP) or long period (LP) was calculated by dividing the number of SP (Period < SP cutoff) or LP mutants (Period > LP cutoff) by the total number of high amplitude filter-passing mutants at that position. These values were color-mapped onto the protein structures, as shown in Figure 3.

Multiple subsets of residues were defined to compare different parameter distributions, as shown in Figure S8–S11. We used Bio.PDB.PDBParser and Bio.PDB.NeighborSearch in the Biopython library to analyze PDB files of the Kai protein structures. Subunit interfaces and protein interfaces were defined as a set of residues with any heavy atom located within 5Å of the other subunit or protein. For KaiA and KaiC subunit interface, PDB 5C5E (*5*) and PDB 3DVL (*6*) were used. Protein-protein interfaces in the nighttime complex were defined using PDB 5JWR (*7*).

The KaiA subunit interface was calculated using 5C5E by calculating distances between chains A and B. Similarly, the KaiC subunit interface was defined based on chains A and B in 3DVL. The KaiB subunit interface in the ground state tetramer was defined based on the PDB 4KSO crystal structure (*8*)chain A, calculating distances to chain B and chain D.

Residue depth was calculated as distance to the nearest water molecule in silico using the DEPTH server (*9*). If the PDB file contained chains that are not part of the region of interest, modified PDB files with those chains removed were used as input. Residues with depth values >

7Å were defined as core residues and residues with depth values < 7Å were defined as surface residues.

The secondary structure information for each KaiB residue in the ground state and fold-switched state was assigned using the STRIDE algorithm (*10*) on chain A in the NMR structure of fold-switched KaiB 5JYT (*7*) and chain A in the crystal structure of ground-state KaiB PDB 2QKE (*11*). Each residue was assigned one of the following: Coil, Strand, Bridge, Alpha Helix, and Turn. Residues for which the secondary structure group assignment changes between folds were included in the KaiB secondary structure flip subset. Using structures 5JYT and 4KSO, residues that are defined as surface in only one of the folds were defined as the KaiB surface flip subset (see Table S1).

The CII-CI waist region of KaiC was defined as residues that were similarly distant to the core of both the CII and CI domain in the 3DVL structure. Specifically, 4 residues in CII and 4 in CI with highest depth values were chosen to represent the cores. For each residue, mean distances to the CII or CI core residues were calculated. Residues with < 20 Å difference in the two distances were defined as the waist subset.

The positions defined based on *Thermosynechococcus elongatus* structures, such as 5JWR, 5JYT, and 2QKE, were mapped to *Synechococcus elongatus* positions based on the alignment of the sequences. Sequence alignment was performed in python using the Bio package function Align.PairwiseAligner() with the following parameters: match_score = 2.0, mismatch_score = –1.0, open_gap_score = –1.0, extend_gap_score = –1.0.

For residue bulkiness calculations, normalized side chain van der Waals volumes for 20 amino acids were obtained from (*12*). For each residue, the Pearson correlation coefficient between the period values and the side chain volumes were calculated for all mutants with high amplitude oscillations.

**Conservation analysis:**

Using the list of KaiABC homologs found by Schmelling et al. (*13*) we created an alignment for KaiA,B, and C using ProMALs3D (cite). To minimize gaps in the alignment we only considered homologs with length over 250aa for KaiA, under 200aa for KaiB and over 400aa for KaiC. From there we used the pysca toolkit (*14*) to further process the multiple sequence alignment, remove gaps, and calculate the Kullback-Leibler (KL) divergence of each position (unweighted by similarity of other sequences) in the alignment compared to the base distribution of amino acid abundance, defined as:

$$D_{KL} = \sum P(x_i)\, log \frac{P(x_i)}{Q(x_i)}$$

Where *i* indexes all 20 amino acids, $P(x_i)$ is the probability that amino acid $x_i$ is found in the multiple sequence alignment at that position, and $Q(x_i)$ is the probability that amino acid $x_i$ is found in the NCBI refseq non-redundant protein sequence database.

We manually applied a cutoff of $D_{KL} > 2.5$ to define highly conserved positions.

**Supplementary Text**

Mathematical Modeling

A simple model of a nonlinear oscillator capable of showing phase-locking phenomena is a phase oscillator coupled to an external driving cycle (*15, 16*). The oscillator is described by an angle θ moving on a circle with angular velocity $\frac{2\pi}{T}$ where *T* is the natural period. The external cycle is described by phase ϕ moving with angular velocity 2π, so $\phi(t) = 2\pi t$.

If the coupling function *f* depends only on the phase mismatch between the oscillator and the external cycle, the dynamics are:

$$\frac{d\theta}{dt} = \frac{2\pi}{T} - f(\theta - \phi)$$

Because *f* must be 2π-periodic, a simple choice is to use Kuramoto coupling: $f(\theta - \phi) = A\sin(\theta - \phi)$, though other smooth periodic functions will produce qualitatively similar results. This gives a two-parameter model:

$$\frac{d\theta}{dt} = \frac{2\pi}{T} - A\sin(\theta - 2\pi t)$$

where *A* is a coupling constant describing the strength of interaction with the environment and *T* is the natural period of the oscillator.

We numerically integrated this oscillator model by selecting a random initial phase on $[0,2\pi)$, integrating the above differential equation for 2 cycles (mimicking the entrainment protocol in the experiment), nd then evaluating the final oscillator phase θ. In Figure 4C, we randomly drew oscillator periods *T* from a Gaussian distribution with $\mu = 1$ day, $\sigma =$ 6 hours. We drew coupling constants *A* from a Gaussian distribution with $\mu = 3\ \text{day}^{-1}, \sigma = 0.075\ \text{day}^{-1}$, so that relative variation in period is 10x greater than variation in the coupling constant.

In this model, conditions for phase-locking can be determined analytically by solving for fixed points where the phase mismatch between the oscillator and the external cycle is constant. The condition is:

$$\theta - \phi = \arcsin\left(\frac{2\pi}{A}\left(\frac{1}{T} - 1\right)\right)$$

Which has solutions only when the argument of arcsin is less than 1 in absolute magnitude. When $A = 3\ \text{day}^{-1}$, these boundaries are at 16.2 hours and 45.9 hours. The short period boundary is marked on Figure 4C.

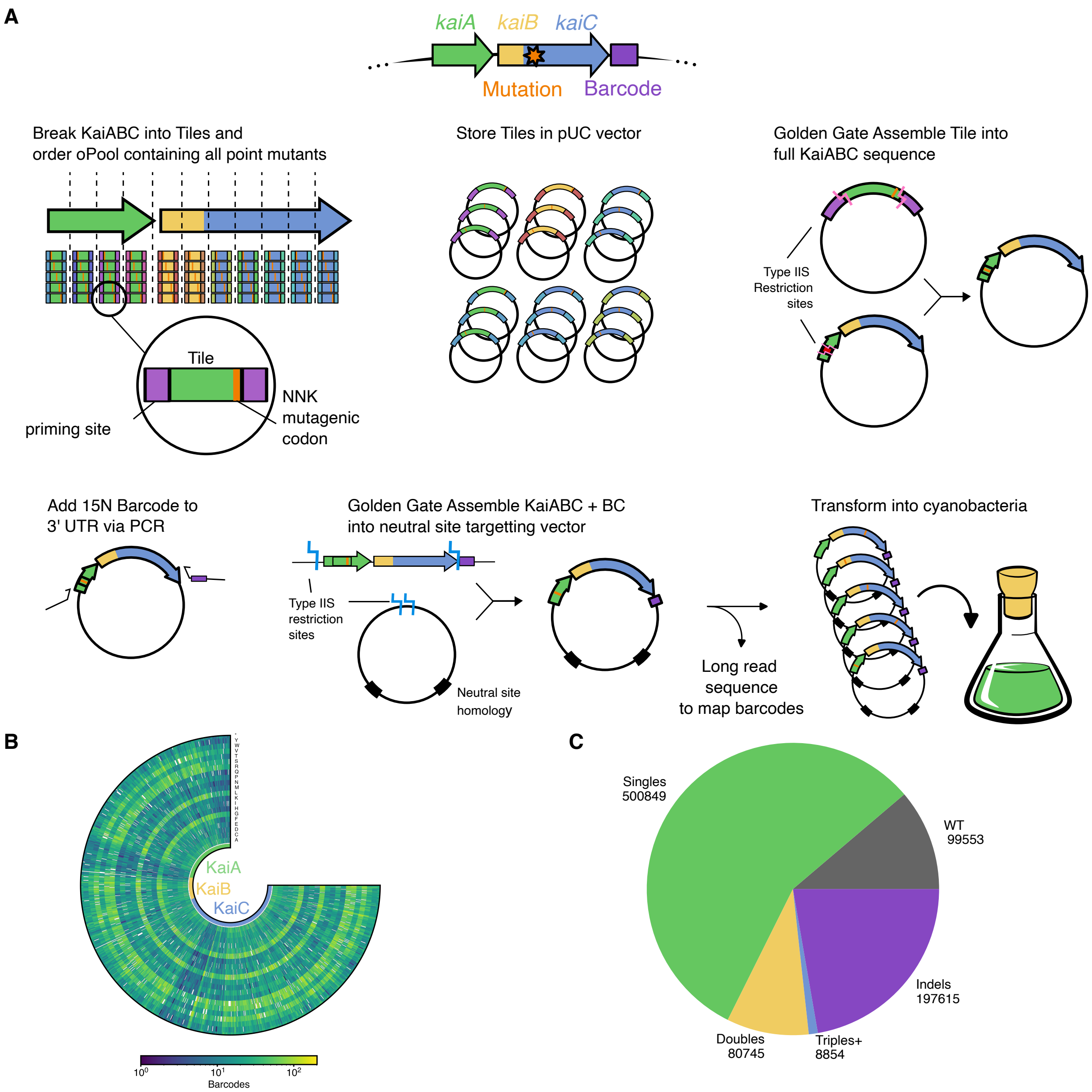


**Figure S1: Creation of a saturation mutagenesis library**

(**A**) Pipeline for library creation. The KaiABC coding sequences were divided into short tiles represented by synthetic pooled oligonucleotides with degenerate NNK sequences at each codon. Each tile was cloned into a small vector for storage, then assembled into the otherwise wildtype kaiABC gene cluster via Golden Gate Assembly. 15N barcodes were added by PCR to the 3' end of *kaiBC* after the STOP codon. The tiles were then combined and cloned into a neutral site targeting vector for cyanobacteria. Long read sequencing was used to map mutations to barcodes. (**B**) Coverage map of single amino acid substitutions in the KaiABC plasmid library. (**C**) Composition of the library by number of mutations and type.

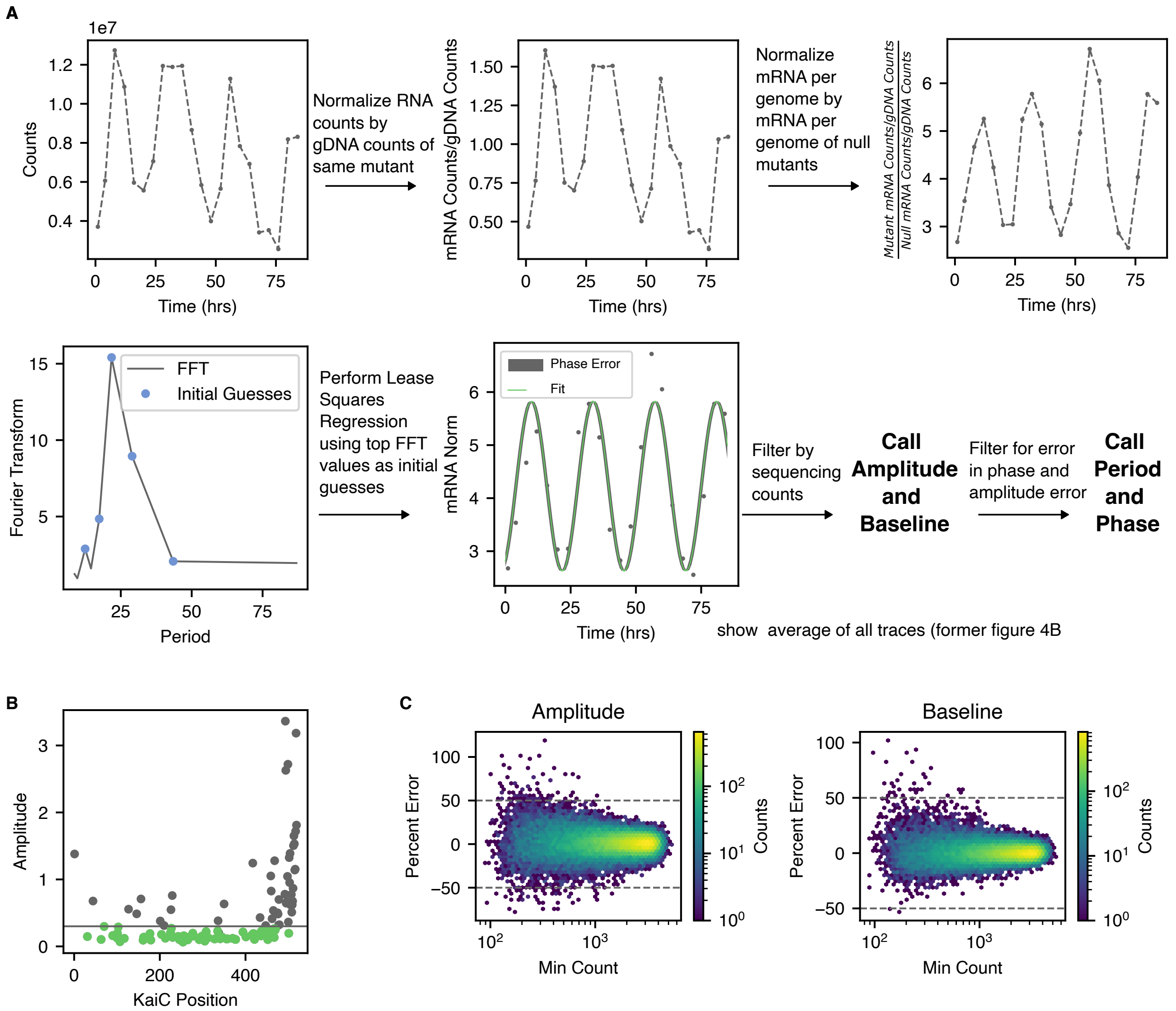


**Figure S2: Fitting and filtering time course data**

(**A**) Analysis pipeline for sequencing time course data. For each timepoint, barcodes mapping to the same mutation were combined then corrected for cell abundance by dividing RNA read counts by gDNA read counts. We then normalize by dividing this signal by RNA counts/gDNA counts for a set of nonsense mutants in KaiC. We analyze mutants that were detected with a minimum number of reads per timepoint. To fit to a sinusoidal model, we first use a fast Fourier transform of the data to get initial period guesses. We then use least squares regression (see Methods) using these initial conditions and pick fit with lowest $\chi^2$. Best fit parameters are used to assign amplitude and baseline. Only fits with sufficiently low estimated error in both phase and amplitude are used to assign a phase and period. (**B**) The best-fit amplitude of all KaiC nonsense mutants. The null normalization set (*green*) was taken as the nonsense mutants with amplitude below the horizontal line (green). (**C**) Error in estimated amplitude (*left*) and baseline (*right*) for subsampled WT barcodes as a function of minimum read count in the time series.

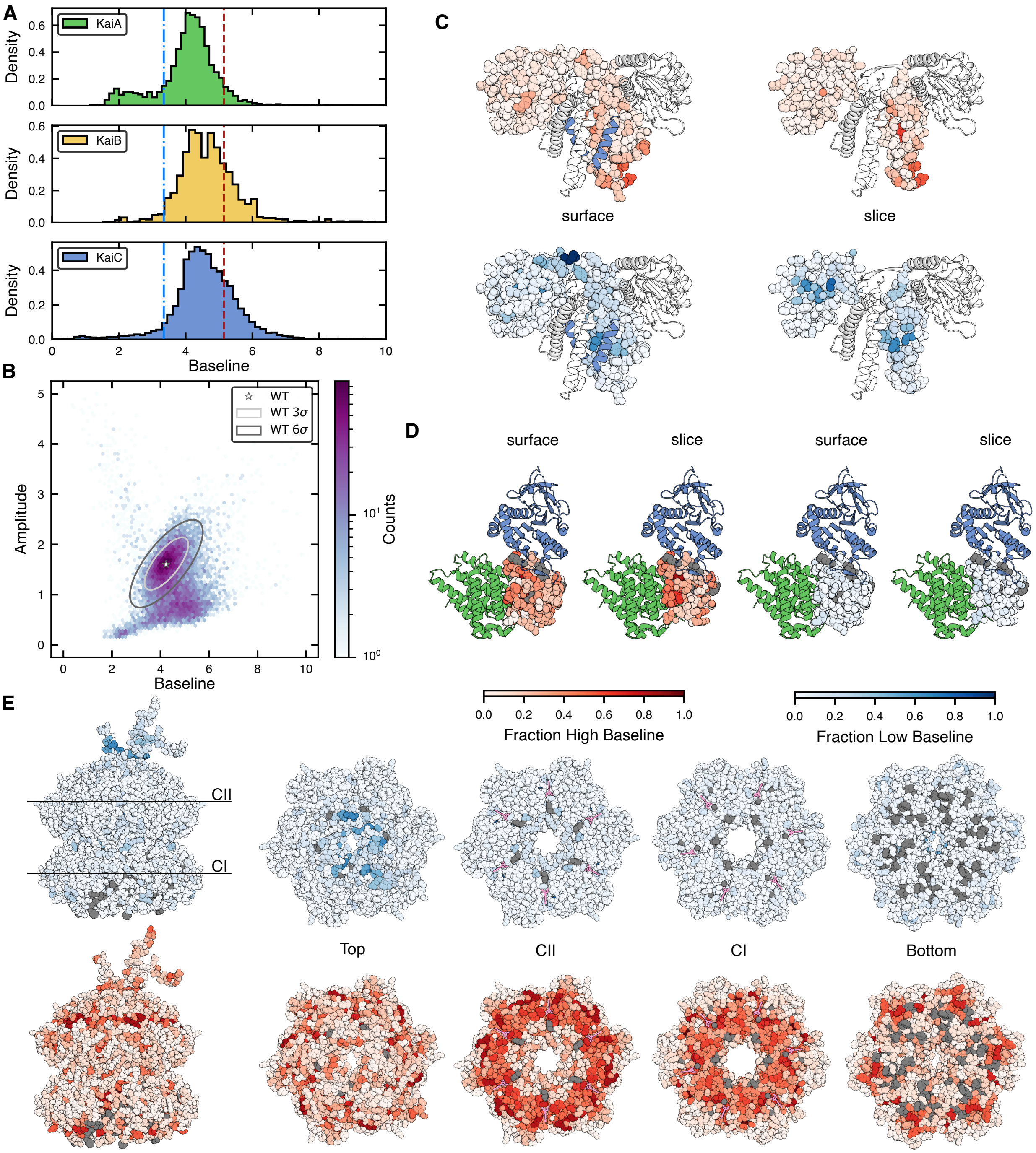


**Figure S3: Mutations affecting baseline**

(**A**) Histograms of baseline in missense mutants by protein. The vertical lines represent 3 standard deviations away from the mean of the WT distribution, these are the cutoffs for high baseline (red) and low baseline (blue). (**B**) Hexbin 2D histogram of baseline vs amplitude. The WT is denoted by a star with ellipses representing the 3rd and 6th standard deviation in WT subsamples. Colorbar is logarithmic. (**C-E**) Fraction low baseline (blue-white color scale) and fraction high baseline (red-white colorscale) shown for KaiC (C, PDB 3dvl), KaiA (D, PDB 5c5e), and KaiB(E, PDB 5jwr). "Slice" indicates that the front half of the protein is hidden to show the protein core.

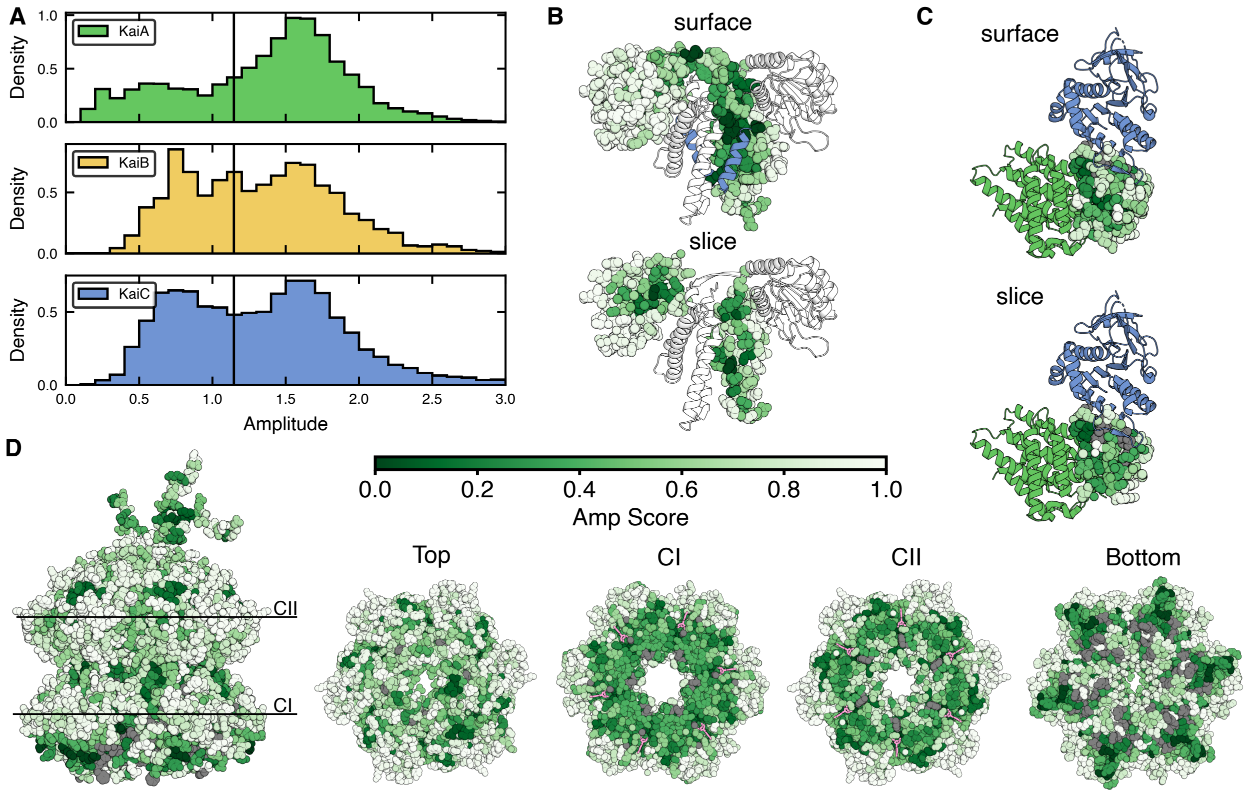


**Figure S4: Additional amplitude views**

(**A**) Histograms of amplitude in missense mutants by protein. The vertical line represents the cutoff between high and low amplitude (defined in Fig. 2). Fraction high amplitude mutants at each position shown using a green-white color scale for (**B**) KaiA (PDB 5c5e). The blue cartoon helix is the KaiC C-terminal peptide. The top structure shows the surface residues of KaiA, and "slice" hides the front half of the protein to show the core. (**C**) KaiB (PDB 5jwr) in the KaiABC complex. KaiC is shown in blue, KaiA is shown in green. (**D**) KaiC (PDB 3dvl), CI and CII views are a top-down view through the indicated plane. ATP is shown in pink.

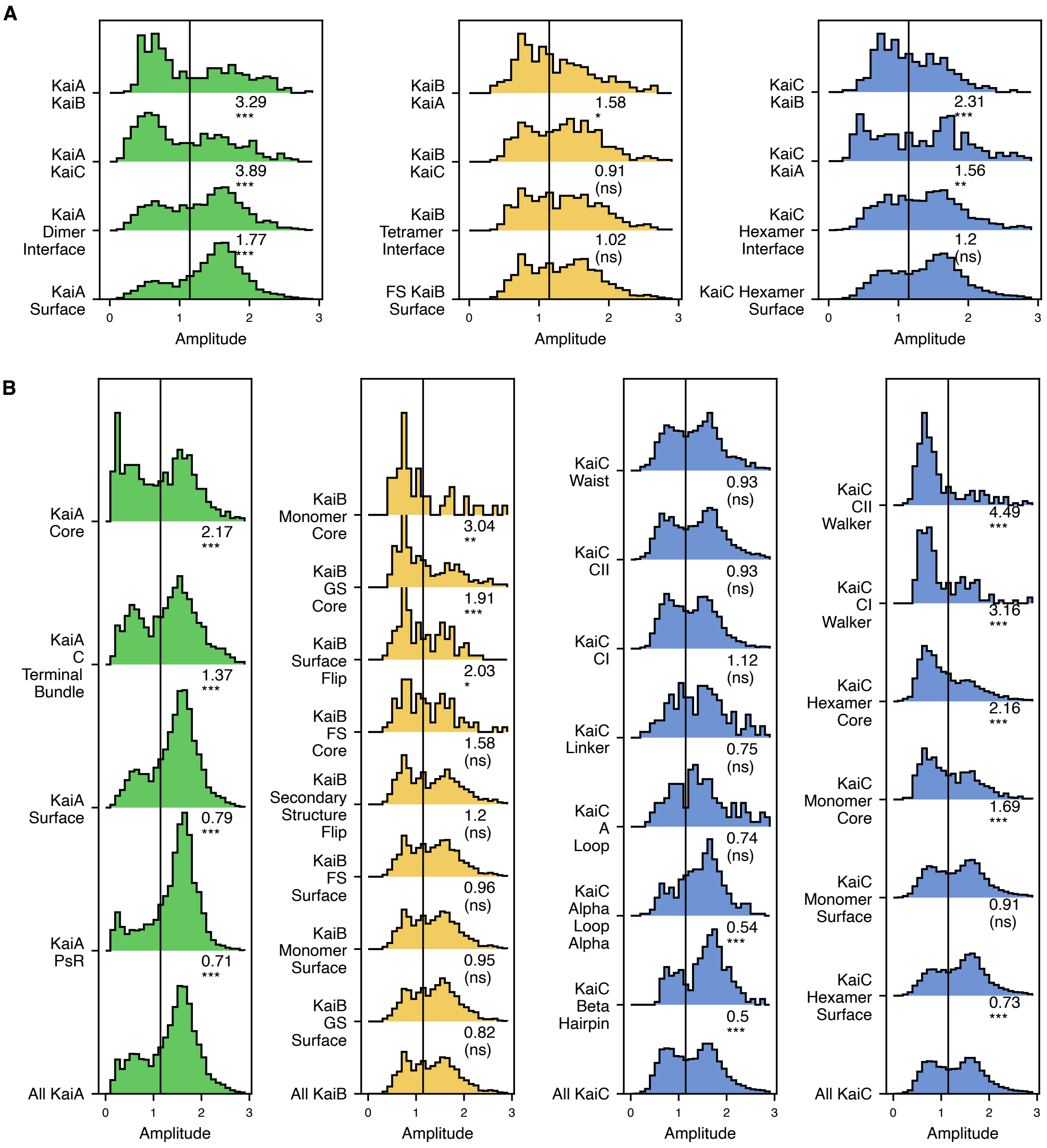


**Figure S5: Distribution of amplitude in structural subsets**

Histograms of amplitudes by subset, and Fisher's exact test of high amplitude mutant counts compared to a reference. See Table S1 for definition of residue subsets. The odds ratio and Bonferroni-corrected p-value are noted under the distribution ($p < 0.001/n$ ***, $p < .01/n$* *, $p < .05/n$ *, $p > .05/n$ (ns) where n is the number of tests (39)). An odds ratio >1 signifies more low amplitude mutants than the reference. (**A**) Protein-protein interfaces tested in reference to the protein surface. (**B**) Subsets tested in reference to the whole protein.

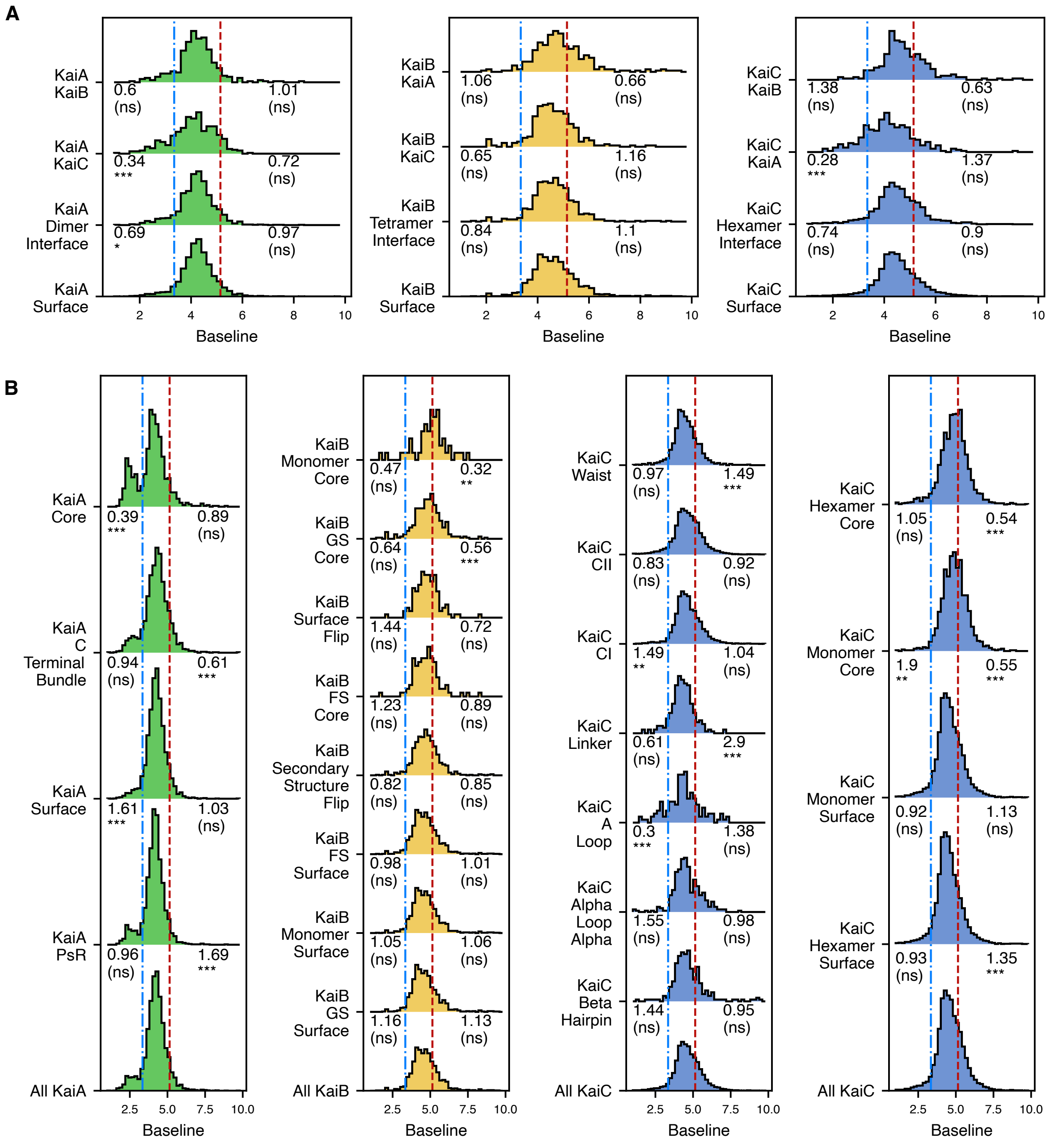


**Figure S6: Distribution of baseline in structural subsets**

Histograms of baseline by subset, the odds ratio by Fisher's exact test for low baseline mutants is shown on the left of the distribution, the high baseline test is shown on the right. See Table S1 for definition of residue subsets. The odds ratio and Bonferroni-corrected p-value are noted under the distribution ($p < 0.001/n$ ***, $p < .01/n$* *, $p < .05/n$ *, p- > .05/n (ns) where n is the number of tests (39)). An odds ratio <1 signifies more baseline mutants than the reference. (**A**) Protein protein interfaces tested in reference to the protein surface. (**B**) Subsets tested in reference to the whole protein.

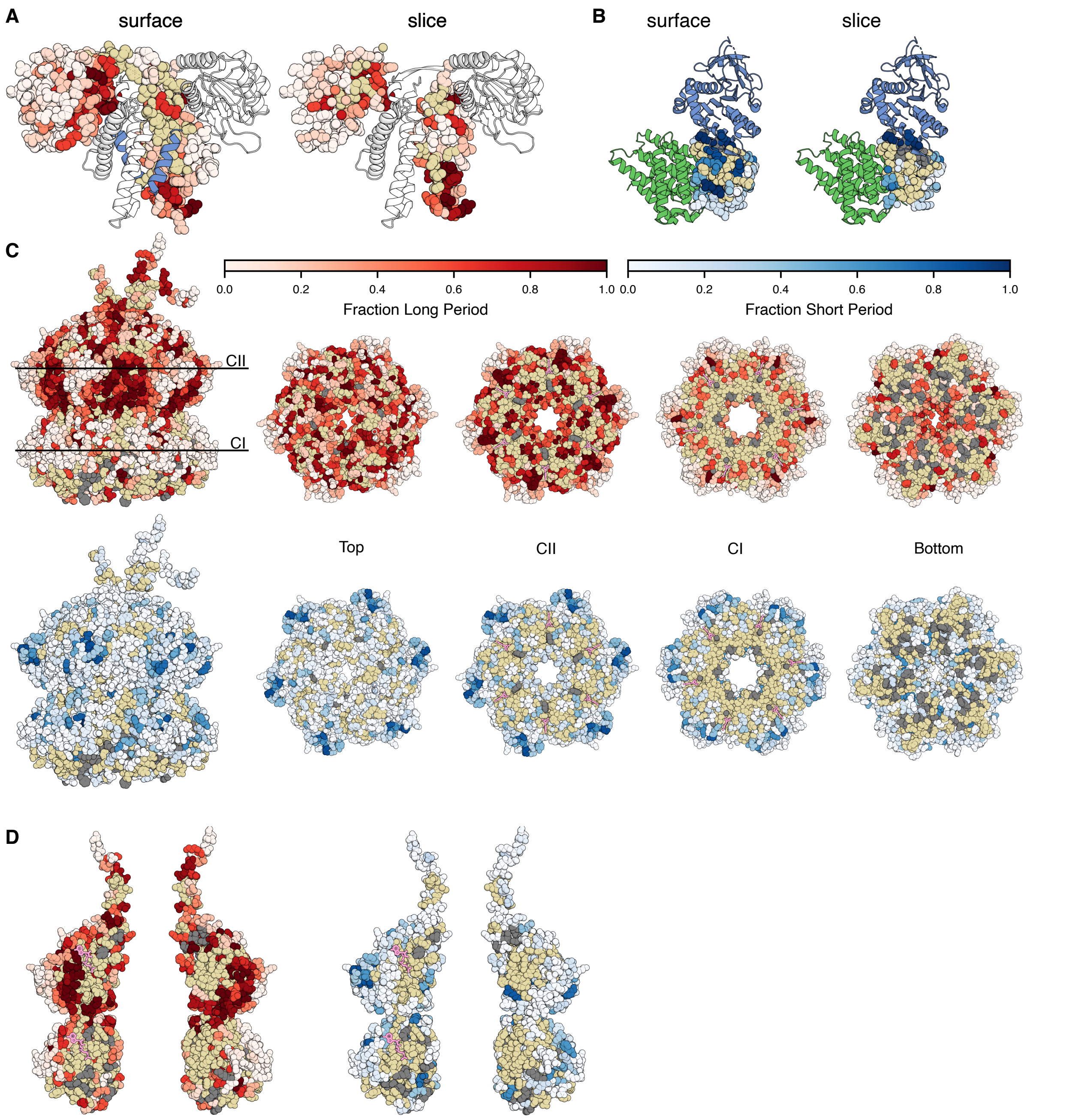


**Figure S7: Additional views of period mutants**

Fraction of short period (red-white color scale) and long period (blue-white color scale) defined as in Figure 3. Beige: fewer than 5 mutants have a defined period. Gray: fewer than 5 mutants present. (**A**) KaiA (PDB 5c5e). The blue cartoon helix is the KaiC C-terminal peptide. The left structure shows the surface residues of KaiA and the "slice" hides the front half of the protein to show the core. (**B**) KaiB (PDB 5jwr) in the nighttime state. KaiC is shown in blue, KaiA is shown in green. The left image shows the surface, the right shows a slice into the core of the protein (**C**) KaiC (PDB 3dvl), CI and CII views are a top-down view through the indicated plane. ATP is shown in pink. (**D**) Views of an isolated KaiC subunit.

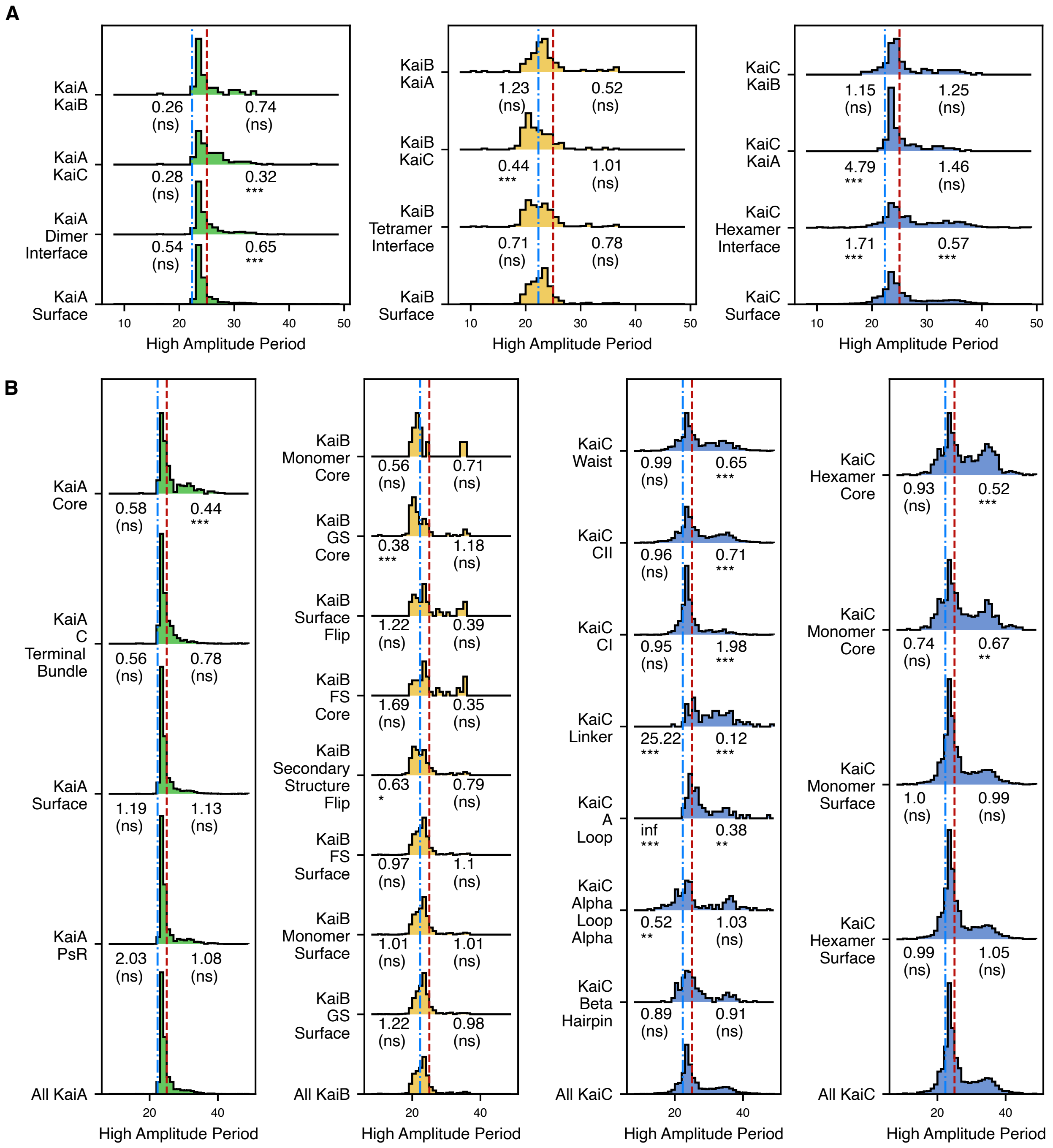


**Figure S8: Distribution of the period of high amplitude oscillations in structural subsets**

Histograms of period of high amplitude mutants by subset, odds ratio, and significance by Fisher's exact test for short period mutants is shown on the left, the long period ratio is shown on the right. See Table S1 for definition of residue subsets. The odds ratio and Bonferroni-corrected p-value are under the distribution distribution (p < 0.001/n ***, p < .01/n* *, p <.05/n *, p > .05/n (ns) where n is the number of tests (39)). An odds ratio <1 signifies more period mutants of that type than the reference. (**A**) Protein-protein interfaces tested in reference to the protein surface. (**B**) Subsets tested in reference to the whole protein.

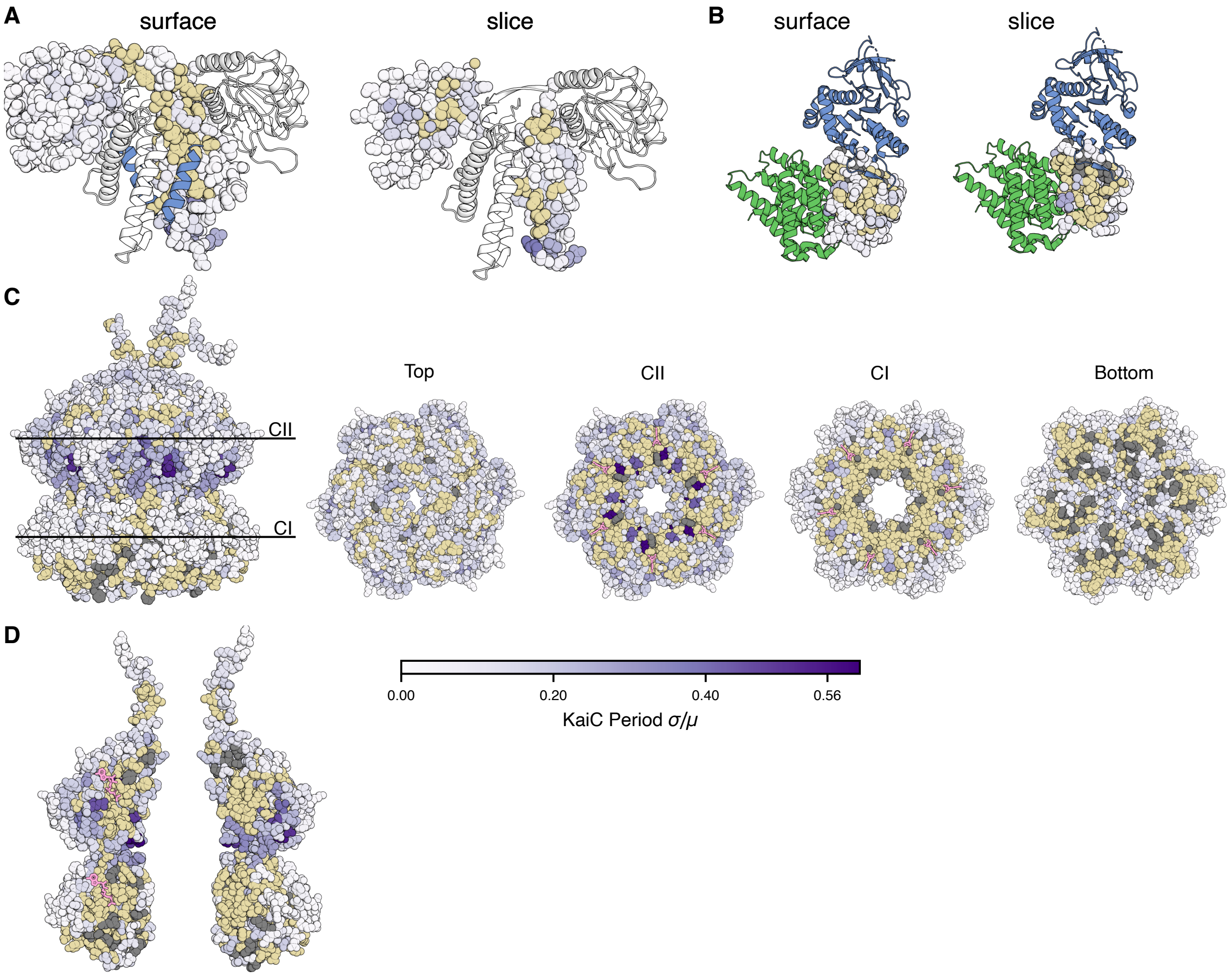


**Figure S9: Additional views of high period sensitivity positions**

Protein structures colored with purple-white color scale to indicate relative standard deviation (standard deviation / mean) of periods achieved by mutation at that position. Beige: fewer than 5 mutants have a defined period. Gray: fewer than 5 mutants present. (**A**) KaiA (PDB 5c5e). The blue cartoon helix is the KaiC C-terminal peptide. The left structure shows the surface residues of KaiA and the "slice" hides the front half of the protein to show the core. (**B**) KaiB (PDB 5jwr) in the nighttime state. KaiC is shown in blue, KaiA is shown in green. The left image shows the surface, the right shows a slice into the core of the protein. (**C**) KaiC (PDB 3dvl), CI and CII views are a top-down view through the indicated plane. ATP is shown in pink. (**D**) Views of an isolated KaiC subunit.

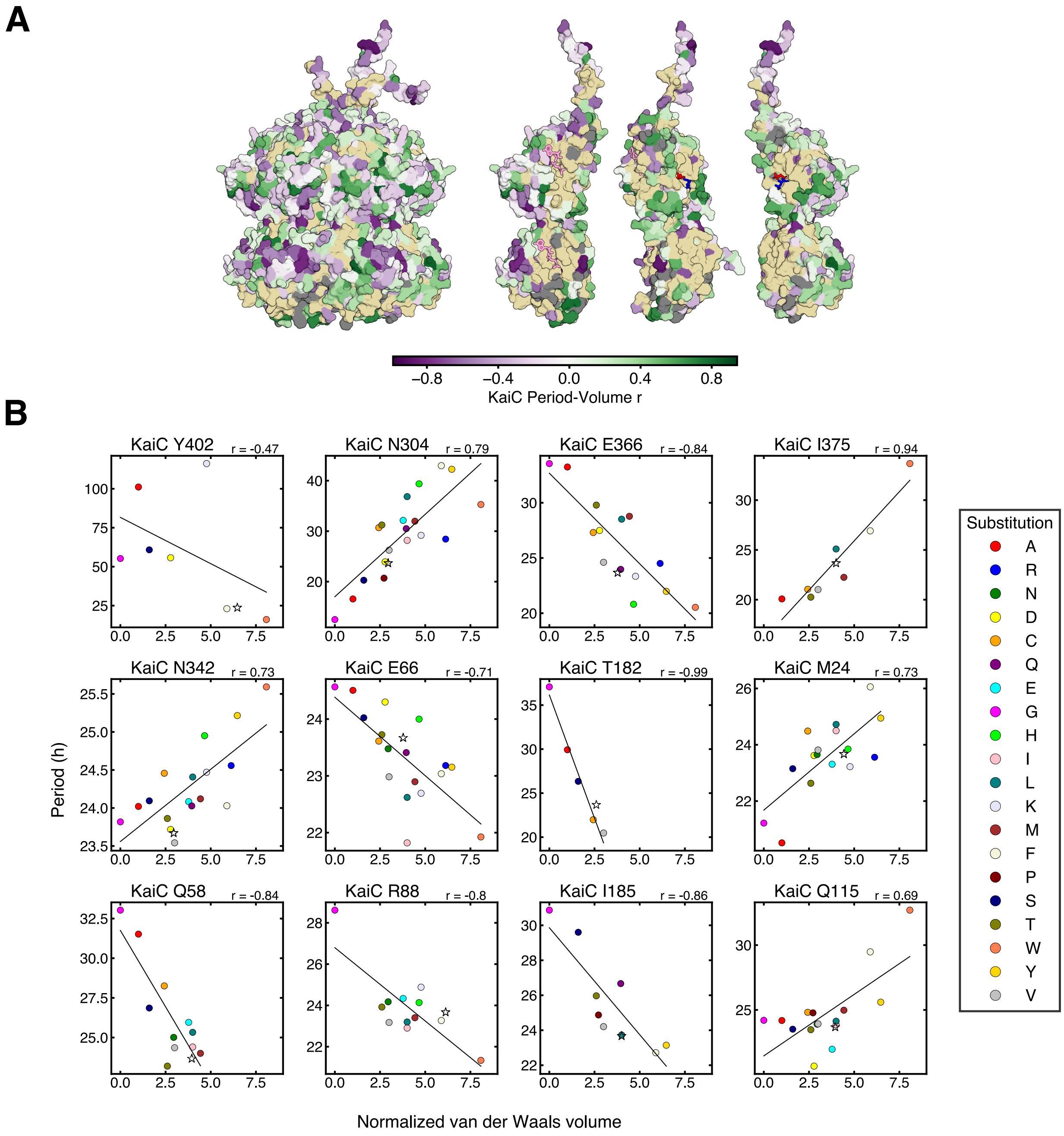


**Figure S10: Correlations and anti-correlations between period and residue bulkiness**

(**A**) KaiC structure where each position is colored by Pearson's correlation coefficient (*r*) between the period and the normalized van der Waals volume of the substituted amino acid for KaiC hexamer (*left*) and isolated KaiC subunits (*right*). ATP is shown in pink. (**B**) Examples of the relationship between the period and bulkiness of the substituted amino acid at a position. Anti-correlation at KaiC Y402, as previously reported by Ito-Miwa *et al.* (*17*) is shown in top left, followed by 11 positions where the correlation has the smallest p-values.

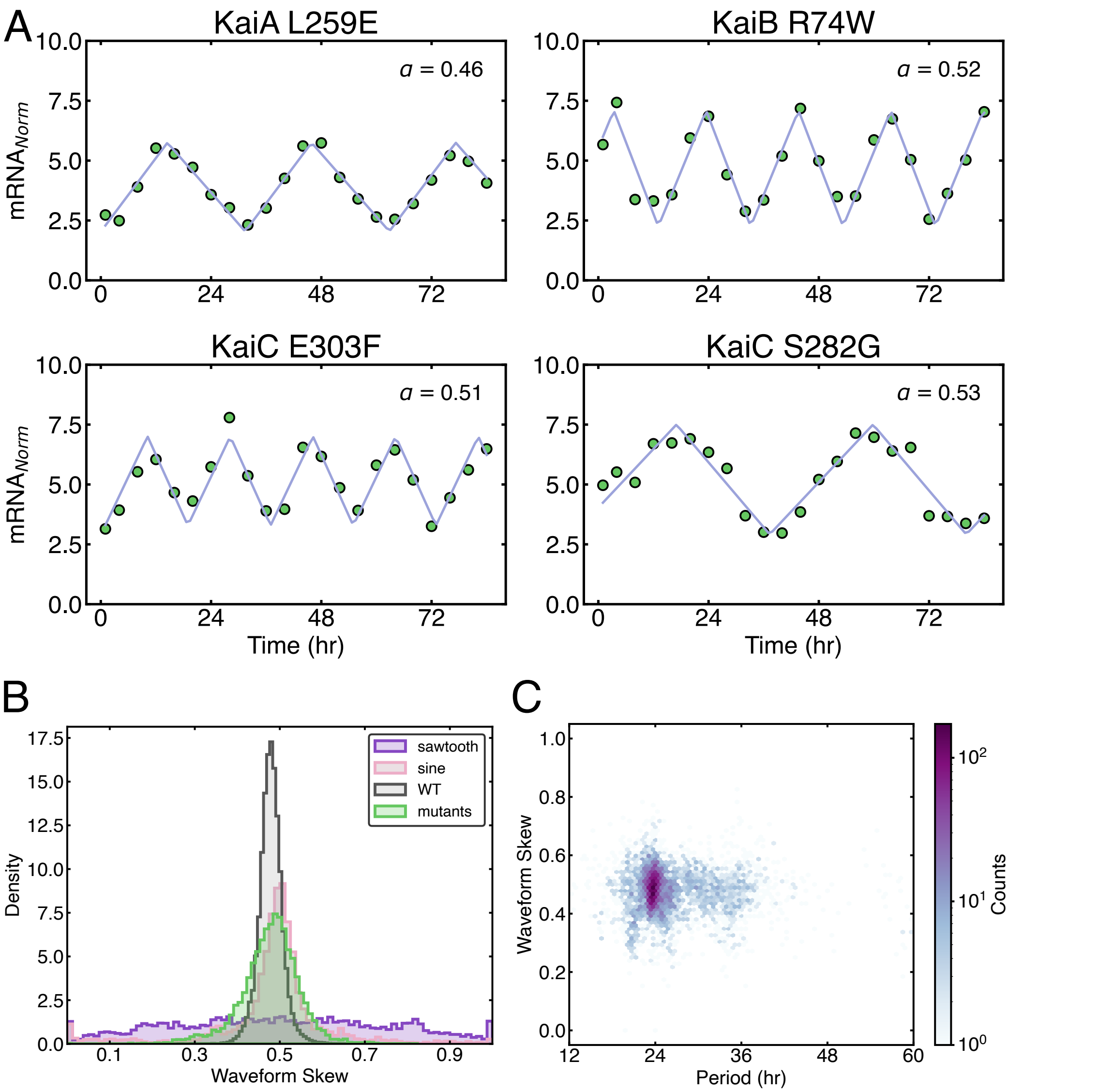


**Figure S11: Waveform shape analysis of mutants**

(**A**) Least-squares regression of a triangle wave model to selected mutants. Waveform skew (α) is a number between 0 and 1 extracted from the triangle wave fit, defined as the time from trough to peak divided by period. $mRNA_{Norm}$ is defined as in Figure 1. (**B**) Distribution of waveform skewness for subsampled WT (*gray*, n=46,875), mutants from the high amplitude group (*green*, n=6,584), and simulated data from sine waves (*pink*, n=7,999) or sawtooth waves (*purple*, n=7,990). Sine waves and sawtooth waves were generated by randomly drawing period, amplitude, and baseline from the mutant phenotypes, then adding 5% normally distributed error to each point. Simulated sawtooth waves have a uniform distribution of skewness. (**C**) Hexbin 2D histogram of waveform skewness vs. oscillator period in mutants from the high amplitude group, derived from triangle wave regression (n=6,584). Color bar is logarithmic.

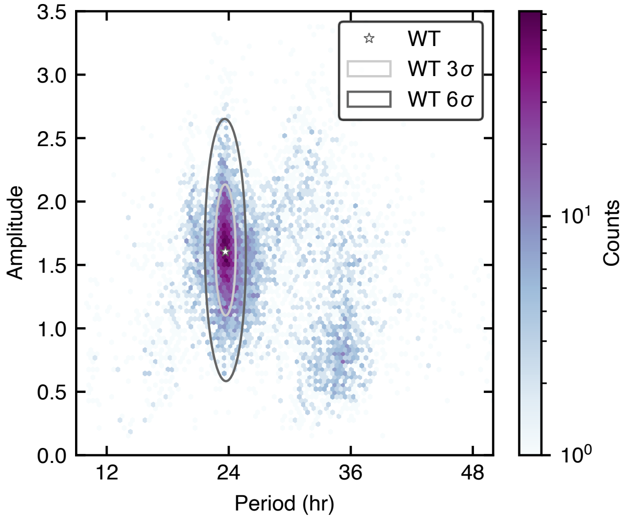


**Figure S12: Dependence of period on amplitude for KaiABC mutants**

Hexbin 2D histogram of amplitude and period for missense mutants passing read count and error filters. Ellipses show 3-sigma and 6-sigma boundaries on fit parameters for subsampled WT barcodes. Color bar is logarithmic.

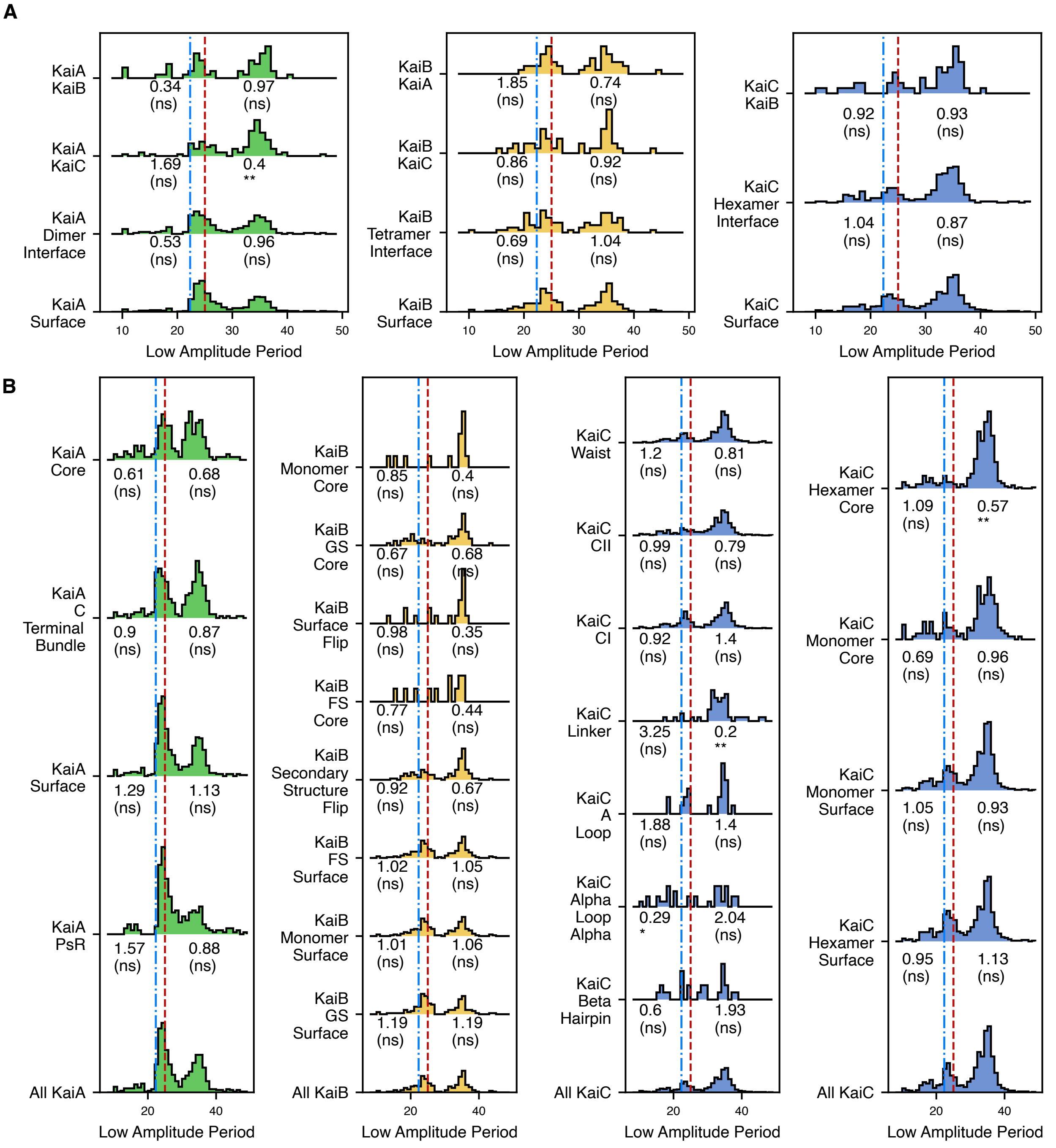


**Figure S13: Distribution of the period of low amplitude oscillations in structural subsets** Histograms of period of low amplitude mutants by subset, the odds ratio of the fisher's exact test for short period mutants is shown on the left, the long period ratio is shown on the right. See Table S1 for definition of residue subsets. The odds ratio and Bonferroni-corrected p-value are noted under the distribution (p < 0.001/n ***, p < .01/n* *, p <.05/n *, p > .05/n (ns) where n is the number of tests (39)). An odds ratio <1 signifies more period mutants than the reference. (**A**) Protein-protein interfaces tested in reference to the protein surface. (**B**) Subsets tested in reference to the whole protein.

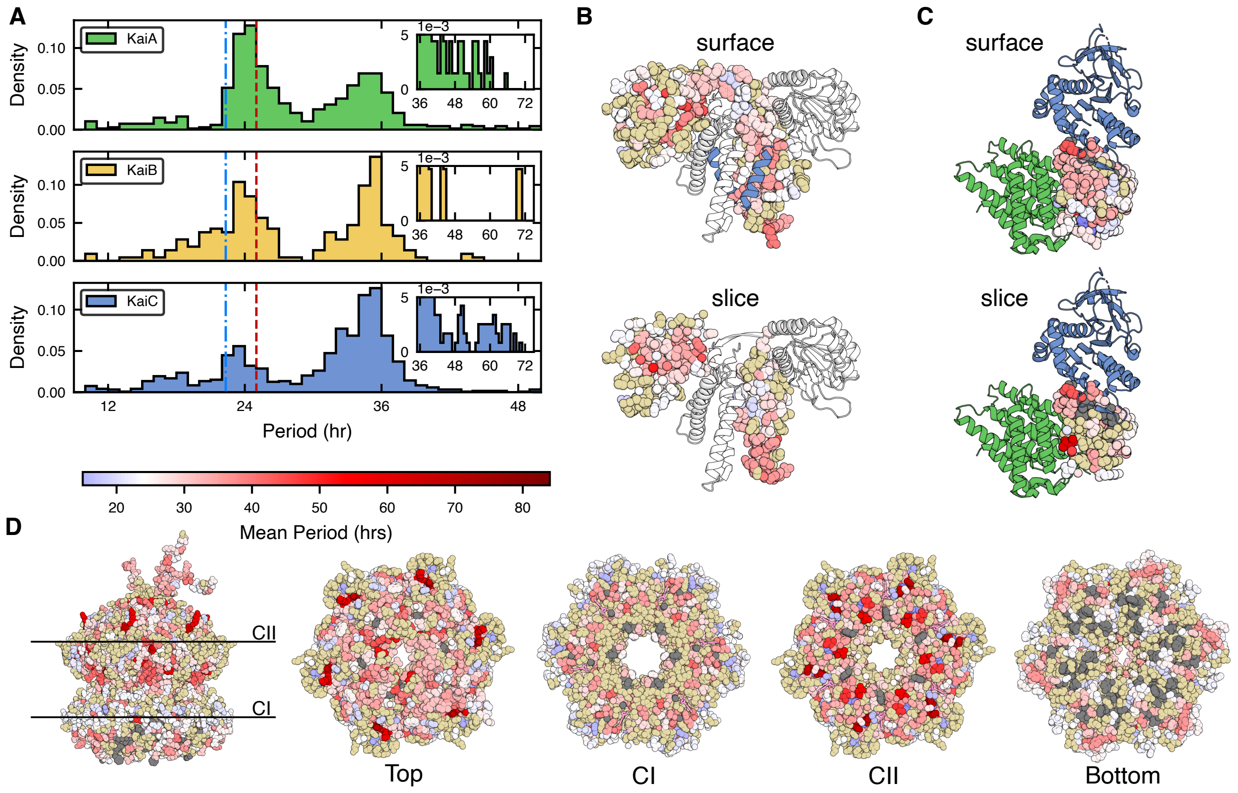


**Figure S14: Period distribution and structural views of low amplitude mutants**

(**A**) Histograms of periods of low amplitude missense mutants by protein. The vertical lines represent the cutoffs for long period (*red dashed line*) and short period (*blue dashed line*). (**B-D**) Structural views colored on a blue-white-red color scale by mean period for low amplitude mutants. Beige indicates no such mutants at that position. (**B**) KaiA (PDB 5C5E). The blue cartoon helix is the KaiC C-terminal peptide. The top structure shows the surface residues of KaiA, and the "slice" hides the front half of the protein to show the core. (**C**) KaiB (PDB 5JWR) in the nighttime state. KaiC is shown in blue, KaiA is shown in green. (**D**) KaiC (PDB 3DVL), CI and CII views are a top-down view through the indicated slices. ATP is shown in pink.

| Structural Element | Residue Numbers |
|---|---|
| KaiC Linker | 248-261 |
| KaiC A Loop | 486-498 |
| KaiC Beta Hairpin | 468-483 |
| KaiC Alpha Loop Alpha | 321-342 |
| KaiC CI | 1-247 |
| KaiC CII | 262-519 |
| KaiC Waist | 24-34,49-51,66,86-93,186-189,207-221,229-257,352-362,364-366,368-369,386-395,397-398,402,405 |
| KaiC CI Walker | 46-53,77-78,151-155 |
| KaiC CII Walker | 288-295,318-319,374-378 |
| KaiA PsR | 1-135 |
| KaiA C Terminal Bundle | 180-284 |
| KaiA Surface | 1-6,11-39,41-50,52,57-69,71-74,82-98,101-113,115-116,123-124,126-159,161-193,195-197,199-200,202-217,227-232,234-235,237-263,265-267,270-271,274-275,277-284 |
| KaiA Monomer Surface | 1-6,11-39,41-50,57-69,71-74,81-98,101-117,119-124,126-132,134-152,154-197,199-200,202-218,220-232,234-235,237-267,269-284 |
| KaiA Core | 7-10,40,51,53-56,67,70,75-83,99-100,106,114,116-122,125,133,160,164,173,178,194,198,201,218-226,228,233,236,264,268-269,272-273,276 |
| KaiA Monomer Core | 7-10,40,51-56,70,75-80,99-100,118,125,133,153,198,201,219,233,236,268 |
| KaiC Hexamer Surface | 14-27,29-35,38-40,49,54,58,61-70,76,79-82,84-129,131-140,150-159,162-163,165-166,169-170,172-177,184-196,209-217,226-232,234,238-269,271-277,279-281,300,303-311,320-322,325-334,336-359,361-362,364-374,381-382,384-394,401-402,405-408,415-418,420-424,426-429,445-451,460-466,468-469,471-488,490-519 |
| KaiC Hexamer Core | 23,28,36-37,41-57,59-60,63,71-75,77-78,83,103,130,141-149,151,160-161,164-165,167-168,171,177-184,191,197-209,218-226,233-237,244,246,270,278,282-299,301-302,312-320,323-324,327,335,350-351,356-357,360,363,375-380,382-383,394-401,403-404,409-414,419,425,430,433-444,452-459,467,470,489,494 |
| KaiC Monomer Surface | 14-22,24-27,29-35,38-40,46-54,58,61-62,64-70,76-102,104-129,131-140,145-146,148-150,152-163,165-166,168-170,172-177,183-196,198-202,204,208-219,221,223-236,238-269,271-277,279-281,289-296,300,303-311,318-323,325-334,336-355,357-359,361-362,364-373,379,381-382,384-394,397-398,400-402,404-408,415-430,433-435,437,442,444-451,454-519 |
| KaiC Monomer Core | 23,28,36-37,41-45,55-57,59-60,63,71-75,103,130,141-144,147,151,164,167,171,178-182,197,203,205-207,220,222,237,270,278,282-288,297-299,301-302,312-316,324,335,356,360,363,374-378,380,383,395-396,399,403,409-414,436,438-441,443,452-453 |

**Table S1**. Definitios of KaiC structural elements. Lists of residues used for analysis of phenotype enrichment.

| Structural Element | Residue Numbers |
|---|---|
| KaiB FS Core | 7,9,11-12,23,27,64-66,85,89 |
| KaiB FS Surface | 1-6,8,10,13-22,24-26,28-63,67-84,86-88,90-102 |
| KaiB GS Surface | 1-6,8,10,13-38,40-56,61-62,65-66,70-71,73-85,92-102,1001-1008,1010,1014-1056,1060-1063,1065-1085,1093-1102 |
| KaiB GS Core | 7,9-13,27,39,46,57-60,63-64,67-69,72,79,86-91,93,1009-1013,1031,1046,1057-1060,1064,1079,1086-1092 |
| KaiB Monomer Surface | 1-8,12-71,73-88,90-102 |
| KaiB Monomer Core | 9-11,72,89 |
| KaiB Secondary Structure Flip | 4-6,14-17,50-79,82-97 |
| KaiB Surface Flip | 7,10,12,23,27,64-66,72,85 |
| KaiA Dimer Interface | 1-3,5,28-29,92-96,103-107,110,112-113,115-120,122-124,126-127,161,163-175,177-178,180-182,186,189-190,192-193,197,200,211,214,217-218,221-222,224-231,234,256,259-263,265-266,269-270,272-284 |
| KaiB Tetramer Interface | 14,34-39,46-47,50-54,56-63,66-70,73-74,79-81,84-88,93,95-97 |
| KaiC Hexamer Interface | 14-18,40,46-49,52,77-78,82,85-86,88-89,109-110,112-114,116,148-149,152-154,157-158,161-162,165-166,169-170,172-173,183-185,188-190,193,195-196,198-200,202,204,206,208-209,211,213-219,221,223-230,232-236,239,250,252-260,279,281,290-291,316-323,326-327,330-331,348-350,352-353,379,381-382,385-386,390,393-394,397,401,404,415,417-426,429,431-433,435,437,439,442,444,446-449,451,454,456-463,465-467,482-491,493-497,499,504,507 |
| KaiA interface with KaiC | 124,127-128,205-206,213,217,220,231,234-235,237-238,241-242,245,255,258-260,262-263,266-267,270-271,274-275,278 |
| KaiC interface with KaiA | 500-503,505-515,517-519 |
| KaiB interface with KaiC | 14,19,46-47,50-51,53-54,57-62,72-81,84,87 |
| KaiC interface with KaiB | 76,104,106-109,111,113-126,128-130,132-133,136,154 |
| KaiB interface withKaiA | 5-8,13,16,24,28,37-45,48-52,55-56 |

**Table S2**. Definitions of KaiA and KaiB structural elements. Lists of residues used for analysis of phenotype enrichment.

**Data S1. RNA sequencing time course and phenotype assignments (separate file)**

period — estimated oscillation period (hours) from least-squares regression
period_err — estimated error in oscillation period (hours) from least-squares regression
baseline — estimated baseline (normalized mRNA) from least-squares regression
baseline_err — estimated error in baseline (normalized mRNA) from least-squares regression
amplitude— estimated amplitude(normalized mRNA) from least-squares regression
amplitude_err — estimated error in amplitude(normalized mRNA) from least-squares regression
phase — estimated phase (radians) from least-squares regression
phase_err — estimated error in phase (radians) from least-squares regression
reduced chi-squared — $\chi^2$ per degree of freedom
Total RNA Reads — total number of jackpot-corrected reads for this mutant
Min Count — number of jackpot-corrected reads at least represented time point for this mutant
Filter Passing — does mutant pass error threshold from least-squares regression?
Amp Group — which amplitude group (defined in Fig. 2)
Base Group — which baseline group (defined in Fig. S3)
Per Group — which period group (defined in Fig. 3)
T01-T22 — normalized mRNA sequencing data
num_muts — number of mutated codons (including synonymous mutations) pooled
codon_muts — list of DNA mutations associated with each barcode pooled for this mutant

**Data S2. Phenotype summary statistics of mutations at each position (separate file)**

High Amp Score — fraction of mutants in the high amplitude group (defined in Fig. 2)
Low Amp Score — fraction of mutants in the low amplitude group (defined in Fig. 2)
Null Amp Score — fraction of mutants in the null amplitude group (defined in Fig. 2)
Mean Period HA — mean period for filter-passing mutants in the high amplitude group
Mean Period LA — mean period for filter-passing mutants in the low amplitude group
LP Score HA — fraction of high amplitude, filter-passing mutants with long period (defined in Fig. 3)
SP Score HA — fraction of high amplitude, filter-passing mutants with short period (defined in Fig. 3)
LP Score LA — fraction of low amplitude, filter-passing mutants with long period (defined in Fig. 3)
SP Score LA — fraction of low amplitude, filter-passing mutants with short period (defined in Fig. 3)
HB score — fraction of mutants with high baseline (defined in Fig. S3)
LB score — fraction of mutants with low baseline (defined in Fig. S3)
Period CV — ratio of standard deviation to mean for periods of high amplitude mutants
KL Divergence — Kullback-Leibler divergence at this position
Number High Coverage Mutants — number of mutants analyzed
Number Error Filter Passing Mutants — number of mutants passing error threshold from least-squares regression

### References


1. S. K. Subramanian, W. P. Russ, R. Ranganathan, A set of experimentally validated, mutually orthogonal primers for combinatorially specifying genetic components. *Synth Biol (Oxf)* **3**, ysx008 (2018).
2. B. Andrews, R. Ranganathan, BCAR: A fast and general barcode-sequence mapper for correcting sequencing errors. *bioRxiv*, (2026).
3. T. Magoc, S. L. Salzberg, FLASH: fast length adjustment of short reads to improve genome assemblies. *Bioinformatics* **27**, 2957–2963 (2011).
4. M. Martin, Cutadapt removes adapter sequences from high-throughput sequencing reads. *EMBnet. journal* **17**, 10–12 (2011).
5. R. Pattanayek, M. Egli, Protein-Protein Interactions in the Cyanobacterial Circadian Clock: Structure of KaiA Dimer in Complex with C-Terminal KaiC Peptides at 2.8 A Resolution. *Biochemistry* **54**, 4575–4578 (2015).
6. R. Pattanayek *et al.*, Visualizing a circadian clock protein: crystal structure of KaiC and functional insights. *Mol Cell* **15**, 375–388 (2004).
7. R. Tseng *et al.*, Structural basis of the day-night transition in a bacterial circadian clock. *Science* **355**, 1174–1180 (2017).
8. S. A. Villarreal *et al.*, CryoEM and molecular dynamics of the circadian KaiB-KaiC complex indicates that KaiB monomers interact with KaiC and block ATP binding clefts. *J Mol Biol* **425**, 3311–3324 (2013).
9. K. P. Tan, R. Varadarajan, M. S. Madhusudhan, DEPTH: a web server to compute depth and predict small-molecule binding cavities in proteins. *Nucleic Acids Res* **39**, W242–248 (2011).
10. D. Frishman, P. Argos, Knowledge-based protein secondary structure assignment. *Proteins* **23**, 566–579 (1995).
11. R. Pattanayek *et al.*, Structural model of the circadian clock KaiB-KaiC complex and mechanism for modulation of KaiC phosphorylation. *EMBO J* **27**, 1767–1778 (2008).
12. J. Meiler, Michael Müller, Anita Zeidler, and Felix Schmäschke, Generation and evaluation of dimension-reduced amino acid parameter representations by artificial neural networks. *Molecular modeling annual* **7**, 360–369 (2001).
13. N. M. Schmelling *et al.*, Minimal tool set for a prokaryotic circadian clock. *BMC Evol Biol* **17**, 169 (2017).
14. O. Rivoire, K. A. Reynolds, R. Ranganathan, Evolution-Based Functional Decomposition of Proteins. *PLoS Comput Biol* **12**, e1004817 (2016).
15. S. Strogatz, *Nonlinear dynamics and chaos : with applications to physics, biology, chemistry, and engineering*. (CRC Press, Boca Raton, ed. Third edition., 2024), pp. pages cm.
16. G. B. Ermentrout, J. Rinzel, Beyond a pacemaker's entrainment limit: phase walk-through. *Am J Physiol* **246**, R102–106 (1984).
17. K. Ito-Miwa, Y. Furuike, S. Akiyama, T. Kondo, Tuning the circadian period of cyanobacteria up to 6.6 days by the single amino acid substitutions in KaiC. *Proceedings of the National Academy of Sciences* **117**, 20926–20931 (2020).
